# Heated debate: Is the seasonal polyphenism of *Araschnia levana* (Lepidoptera: Nymphalidae) driven by thermoregulation?

**DOI:** 10.64898/2026.02.08.704656

**Authors:** Daniel Linke, Jan Okrouhlik, Alena Sucháčková Bartoňová, Leonardo Ré Jorge, Pável Matos-Maraví, Irena Klečková

## Abstract

The seasonal forms of the temperate butterfly *Araschnia levana* (Nymphalidae: Nymphalinae) differ in morphology (weight, wing area, and wing loading) and colouration. Spring individuals are predominantly orange with higher weight per wing area, (i.e. wing loading) while summer individuals are black with a white stripe and have lower wing loading. However, it remains unclear if and how these seasonal differences affect heating and cooling dynamics. We compared thermal responses of seasonal forms, focusing on the roles of morphology and colouration. Further, we assessed whether live butterflies heat and cool differently from dead individuals to detect the presence of active thermoregulation.

Morphology and colouration influenced the thermal dynamics of the thorax and wings as expected from heat-transfer principles, but we found no evidence of active thermoregulation on the thorax. Based on aligned temperature curves, seasonal forms showed similar thermal dynamics. This similarity was driven by morphology and colouration, with larger wing area accelerating thermal change and higher body weight (or wing loading) reducing it, thereby masking underlying form-specific patterns. After accounting for significant morphological differences between forms, the thorax of spring individuals heated and cooled faster than that of summer ones.

This trend suggests form-specific optimisation of thermal performance, likely as a response to temperate climates. Thermal responses differ between forms in ways not directly explained by the polyphenism itself, potentially reflecting a broader trait of multivoltine ectotherms to cope with seasonal temperature changes.

## Introduction

As ectotherms, butterflies rely heavily on external heat sources to regulate their body temperature, making thermoregulation a key trait influencing activity patterns, distribution and individual fitness. The rate at which butterflies heat and cool is influenced by morphology, physiology and behaviour. Morphological traits influencing thermal regulation are complex and include body size, mass, colouration, and cuticular structures (Wang et al., 2021; Markl et al., 2022; Kleckova, et al., 2023a). Physiological mechanisms beyond morphology also influence body temperature regulation in insects. Haemolymph circulation in insect wings and bodies can distribute heat, nutrients, and support sensory hairs and other living tissues (Salcedo & Socha, 2020), and a range of physiological and molecular responses modulate thermal tolerance and body-temperature plasticity across insect taxa (see review: Lahondère, 2023). In butterflies specifically, haemolymph movement plays a role in redistributing thermal energy (Tsai et al., 2020), affecting heating and cooling dynamics. Lastly, behaviour and micro habitat choice, such as adjusting wing positions during basking (Kemp & Krockenberger, 2002), producing endogenous heat via wing vibration (Rosa & Saastamoinen, 2020) or selecting specific microhabitats to heat or cool (Kleckova et al., 2014).

In some multivoltine species, generations have different phenotypes that influence thermoregulation, such as higher heating capabilities during the colder season, which is referred to as polyphenism. For example, the dark red hindwings in the autumn generation of *Junonia coenia* in North America likely increase heating rates (Järvi et al., 2019). Seasonally polyphenic species have evolved in highly seasonal environments, where repeated environmental changes shape life-history strategies (Halali et al., 2024). Because these cycles are predictable, the conditions during the larval stage serve as a reliable predictor of the environment that adults will encounter. In these species, environmental cues experienced during the larval stage trigger the development of distinct adult phenotypes (Nelson et al., 2010). These cues initiate gene expression pathways that drive the development of season-specific adult forms, which are directly exposed to natural selection (Van Dyck & Wiklund, 2002).

A well-known example of such seasonal polyphenism is the map butterfly, *Araschnia levana* (Lepidoptera: Nymphalidae: Nymphalinae), occurring across temperate Europe to East Asia. The development of the species’ seasonal forms is regulated primarily by the photoperiod experienced during the larval stage, indicative of the upcoming season (Nelson et al., 2010; Esperk & Tammaru, 2021). Short-day conditions induce the orange spring form, whereas long-day conditions trigger development into the black summer form. Despite their distinct colouration, a recent field survey showed similar thoracic–ambient temperature relationships in both forms (Laird-Hopkins et al., 2023). The spring form exhibits higher wing loading (i.e., weight per wing area) and brighter orange wings, while the summer form has darker wings with a broader white stripe and a lower wing loading due to larger wings (Figure 1) (Fric & Konvicka, 2002; Nelson et al., 2010; Esperk & Tammaru, 2021). Although larger wings have been associated with increased dispersal capabilities (Sekar, 2012), they typically result in slower thoracic heating rates (Kleckova, et al., 2023b), which may invoke evolutionary trade-offs. While colour differences are the most apparent, seasonal forms also differ in gene regulation of antibacterial activity (Vilcinskas & Vogel, 2016; Baudach et al., 2018).

**Figure 1:**
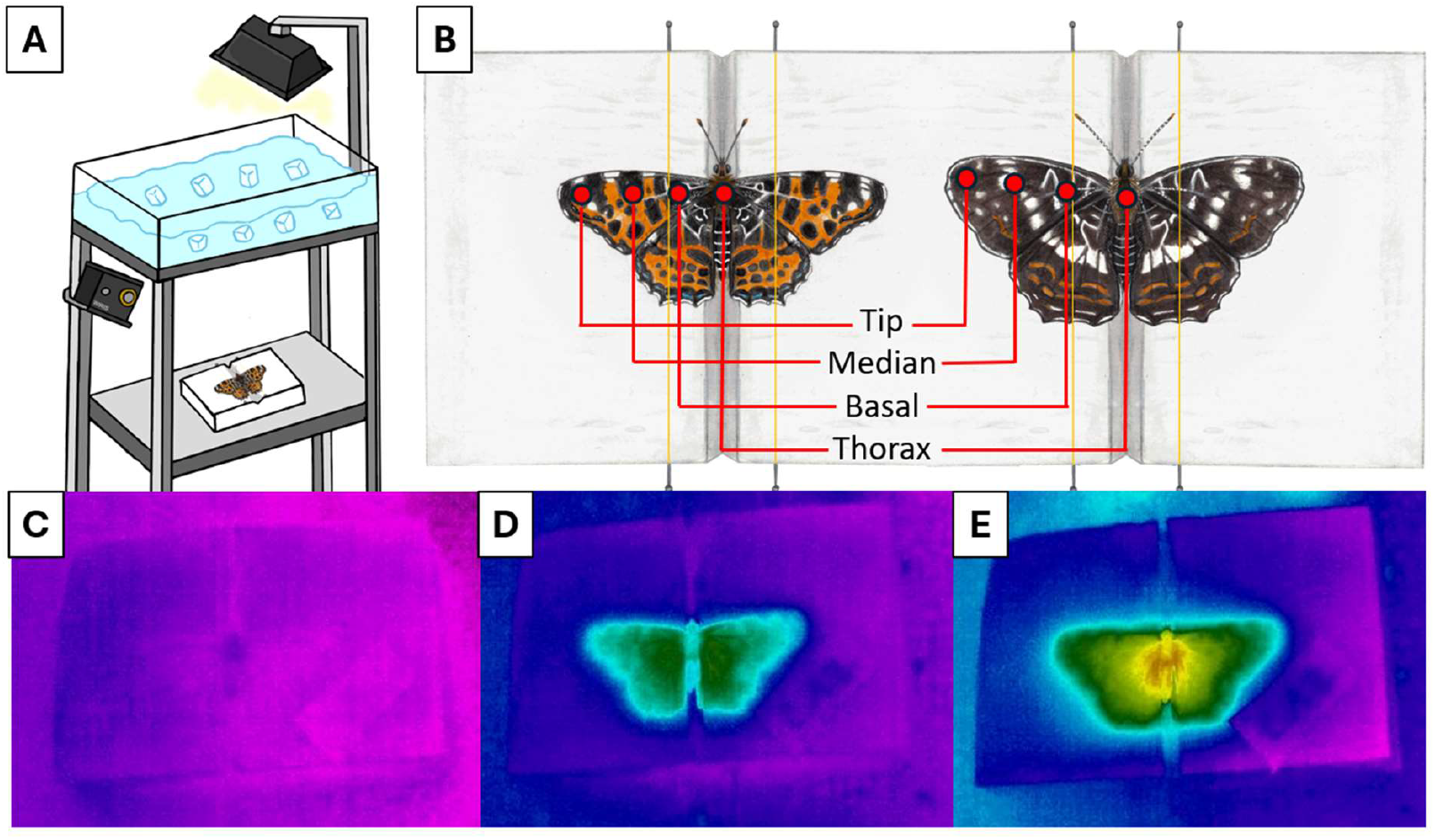
**A**: Thermal camera set up in the climate chamber; **B**: Individuals of *Araschnia levana* were attached by thin blond hairs to the holder in standardised wing position and regions of interest (ROIs) for which temperatures were measured; **C-E**: Examples of *A. levana* during the heating experiment (L012 - Alive - Heating), **C**: Before starting the experiment, **D**: 30 seconds and **E**: 180 seconds of heating. Drawings by Michaela Helclová.

While these morphological and physiological differences between forms point towards different thermal responses, it remains unknown whether these differences translate into measurable variation between heating and cooling dynamics of the seasonal forms of *A. levana*. Understanding this aspect is crucial to uncovering a possible adaptive origin of seasonal polyphenism. Within this manuscript we aim to test the following questions:

1. Do the seasonal forms of *Araschnia levana* differ in their heating and cooling dynamics, and is this variation explained by differences in colouration?
2. Can haemolymph circulation in the wings of *A. levana* modulate thoracic heating or cooling dynamics, indicating active thermoregulatory mechanisms?
3. If variation in heating and cooling dynamics is not solely explained by seasonal form (spring vs. summer) or colouration, which additional factors, i.e., weight, wing area or wing loading, account for thermoregulatory differences?

## Methods

### Butterfly collection

Butterflies were caught using entomological nets in the surroundings of České Budějovice, Třeboň and Mariánské Lázně in Czechia during 2023 (Appendix S1). Live butterflies were stored in glassine envelopes and transported to a temperature-controlled room set to 10°C at the University of South Bohemia in České Budějovice. Heating experiments were conducted within 48 hours following the collection of the butterfly. Recently emerged individuals were preferred (i.e., not showing wing wear), but due to low population densities, slightly worn individuals were also included in this study.

### Experimental setup and thermal imaging

To investigate the heating and cooling dynamics of *A. levana*, we employed infrared thermography to measure surface temperatures using a Workswell WIRIS V2 thermal imaging system (Workswell s.r.o., Prague, Czechia). The camera had a thermal sensitivity of 30 mK, an accuracy of ±2%, and was factory calibrated. Prior to each trial, a non-uniformity correction was performed to calibrate thermal readings. To simulate daylight radiation, we used a halogen incandescent linear light bulb (Ecolite model J500-118, 500W, 3050K, 10,000lm, Wolfhausen, Switzerland), filtered through a 5-cm layer of ice-cold water to minimise infrared interference (Figure 1A). All experiments were conducted in an ∼15m^3^ temperature-controlled room set to 10°C. Prior to each experiment the air mixing fans were running continuously, and the room temperature was allowed to stabilise within 0.5 of the set temperature. To minimise air currents in the room during the experiment, the cooling and the air mixing fans were switched off 30s before the start of the heating phase.

To facilitate handling, live butterflies were cooled on ice before placing them on an experimental holder in a standardised wing position (Figure 1B). To prevent movement, wings were secured using fine blond human hair attached to two metal pins (Järvi et al., 2019). The butterfly was positioned approximately 40 cm below the light source, with the thermal camera perpendicular to it. The experiment consisted of two phases: heating phase – the light source was switched on, and the butterfly’s surface temperatures were recorded for 300 seconds, and cooling phase – the light source was turned off, and temperature recordings continued for an additional 300 seconds. Raw thermograms were taken every 2 seconds (Figure 1C-E). Following, the butterfly was euthanized by exposure to chloroform vapours for a minimum of 5 minutes. The experiment was then repeated with the freshly dead butterfly placed in the standard wing position.

After the experiments, the wet body mass of each butterfly was measured using an electronic laboratory scale (model GR-200, A&D, Tokyo, Japan; precision: 0.1 mg), and the specimens were mounted for further morphological analysis.

### Temperature data extraction from thermograms

We determined temperatures in four specific regions of interest (ROI): tip, median and basal parts of the left forewing, and the thorax (Figure 1B); if the left forewing moved, the right forewing was utilised (only in two specimens). The measured area covered approximately 6 mm^2^ in each body part, which represented 114 pixels of the radiometric image distributed in circular shape. The three measured regions on the wing surface were equidistant.

The raw radiometric data were batch processed using Workswell ThermoFormat software v. 3.4.49.349 (Prague, Czechia) with an emissivity of 0.95 following previous experiments (Tsai et al., 2020). Surface temperatures were calculated from the raw thermographs using the butterfly–camera distance and the initial room humidity and air temperature as input parameters. Temperature data were then exported from the program into a timestamped CSV file. For each time point, the mean temperature of each of the four study regions was then calculated by averaging respective pixel values (further information regarding quality control see Appendix S2).

### Morphometric measurements

Mounted specimens were photographed as described in Linke et al. (2025) with a colour checker card (colorchecker classic mini, Calibrite, Wilmington, USA). Bulk colour scaling was done using RawTherapee v. 5.8 (Budapest, Hungary). Morphometric measurements were taken using GIMP 2.10.24 (GNU Image Manipulation Program) and included forewing length (cm) and the total wing area (sum of all four wing areas, cm^2^). Wing loading (mg cm^-2^) was calculated as the ratio of wet body mass to the total wing area. Wet body mass, measured immediately after the experiments, was preferred over dry weight to account for the variance in water content between specimens (ranging from 60 to 76%). Average specimen darkness/lightness was calculated using colour-corrected photographs and the program ImageJ v. 1.54 (Schneider et al., 2012). For easier use the colour was scaled between 0 and 1, with 1 being fully white any 0 completely black. As some individuals showed slight wing wear, colour measurements may be biased due to specimen age; however, they reflect the colouration present during the experiment.

Using linear models, we compared forewing length, total wing area, wing loading and average colour (responses) with form and sex as categorical predictors. Post hoc differences were calculated using *emmeans* v. 1.11.2-8 (Lenth, 2025).

### Modelling of thermal dynamics

We used two approaches to describe the heating and cooling dynamics of the four ROIs (thorax and the three wing regions), (1) estimating parameters of thermal curves, and (2) using temperature differences through time.

First, we fitted the temperature data through time with an exponential asymptotic model:

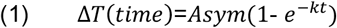

where Asym is the asymptotic temperature and k is the curve shape parameter controlling the steepness of the curve. For each specimen, ROI, and phase, temperature time series were normalised by subtracting the temperature at time 0 (T_0_) from the temperature at each subsequent time point (i.e., T_30_ − T_0_). This aligns curves to a common initial zero and makes the intercept term unnecessary. For cooling curves, temperature values were treated as absolute values prior to fitting, allowing both heating and cooling curves to increase towards Asym, standardising curve direction. Biologically, the parameter k describes how quickly the temperature approaches Asym: higher values of k correspond to faster heating or cooling, whereas lower values indicate slower thermal responsiveness. Starting values for the nonlinear fitting were estimated directly from the data to improve convergence. The asymptotic function was modelled using the *nlme* 3.1-168 package (Pinheiro et al., 2025) in R (R Core Team, 2025).

We compared Asym and k between seasonal forms (spring vs. summer individuals) and specimen status (alive vs. dead) by fitting linear models (*lm*) using the base *stats* package in R; pairwise contrasts were estimated using *emmeans*. Analyses were performed separately for heating and cooling experiments, and for all four ROIs.

In the second approach, we calculated temperature differences (ΔT) of each ROIs at multiple time points, 10, 30, 60, 180 and 300 seconds since experiment start, relative to the baseline temperature, T_0_ (i.e., ΔT_30_ = T_30_ - T_0_). Temperature data through time was smoothed to reduce random noise in the time series, using a locally weighted polynomial regression (LOWESS smoothing) implemented in the R package *stats* v. 4.5.1 (Cleveland, 1979). Butterflies which showed random abrupt changes in the smoothed heating or cooling curves (e.g., butterfly moved its wings during the measurements) were excluded from the analyses.

To assess the effects of morphology (wing loading, weight, wing area), colouration, seasonal form, sex and specimen status on ΔT through time, we used linear mixed-effects models (LMMs) implemented with the *lmer* function from the *lme4* R package v. 1.1-37 (Bates et al., 2015).

Due to wing loading being a derived metric from weight and total wing area we fitted two separate models including either (2a) wing loading or (2b) weight and wing area. All morphometric parameters were scaled to ensure equal variance and comparability.

(2a) Wing loading model:

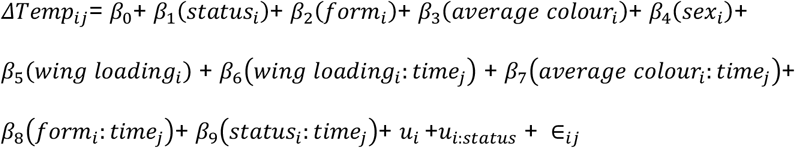

The 2b model includes effects of weight and total wing area and their interaction with time instead of wing loading; the remaining model syntax is identical to model 2a. β_0_ is the fixed intercept, β_1_ to β_n_ are the fixed effect coefficients, time is treated as a factor for the five time intervals, u_i_ and u_i:status_ are random terms for each individual and for each individual × status combination, and ε_ij_ is the residual error. The seasonal form was included separately for heating and cooling, as physiological differences may lead to distinct thermal dynamics independent of colouration. Pairwise contrasts for categorical factors (status and form) were obtained using the *emmeans* package v. 1.11.2-8 (Lenth, 2025).

As starting temperatures were standardised separately for heating and cooling, the asymptotic temperature reached during cooling is influenced by the total heat gained during the preceding heating phase. Thus, any differences in heating (for example due to form-specific morphology or colouration) will affect cooling, especially in the later stages as temperatures approach the ambient asymptote. Early cooling (when body temperature is still far above ambient) accurately reflect cooling dynamics.

All statistical analyses were run in R v. 4.5.0 (R Core Team, 2025), and graphics were generated using ggplot2 v. 3.5.2 (Wickham, 2016).

## Results

### Morphological differences between seasonal forms and sexes

In total we measured 21 spring individuals (18 females and 3 males) and 21 summer individuals (7 females and 14 males). Linear models showed that forewings were significantly shorter in the spring than in the summer form (t_39_ = 6.71, p < 0.001). Regardless of form, females had significantly longer forewings compared to males (t_39_ = −5.72, p < 0.001), the model explained 57% of forewing length variation (R^2^= 0.569). Total wing area followed the same pattern, with summer individuals exhibiting significantly larger wings (t_39_ = 6.74, p < 0.001). Females had significantly larger total wing area than males (t_39_ = −7.90, p < 0.001), with sex and form explaining 65% of total wing area variation (R^2^= 0.646). Wet weights were different between sexes (t_39_ = −3.69, p < 0.001), with females being heavier, but not between forms (t_39_ = 0.56, p = 0.577), explaining 29% of weight variation (R^2^= 0.296). Wing loading was significantly higher (t_39_ = -2.22, p = 0.032) in spring than in summer form but did not differ between sexes (t_39_ = -0.53, p = 0.601), the model explained 19% of wing loading variation (R^2^= 0.188). Thus, spring females had roughly the same size as summer males. The spring form exhibited significantly lighter average colour than the summer form (t_39_ = -3.63, p < 0.001). Across both forms, females had significantly lighter colouration than males (t_39_ = −3.94, p < 0.001), the model explained 61% of total colour variation (R^2^= 0.611). See Appendix S3 Table S2 for average values of morphometric measurements.

### Responses to thermal curve parameters

When comparing the two parameters of the exponential model on the thorax, no differences in the asymptotic temperature (Asym) nor the curve shape parameter (k) were found between seasonal forms or specimen status (Table 1).

**Table 1:**
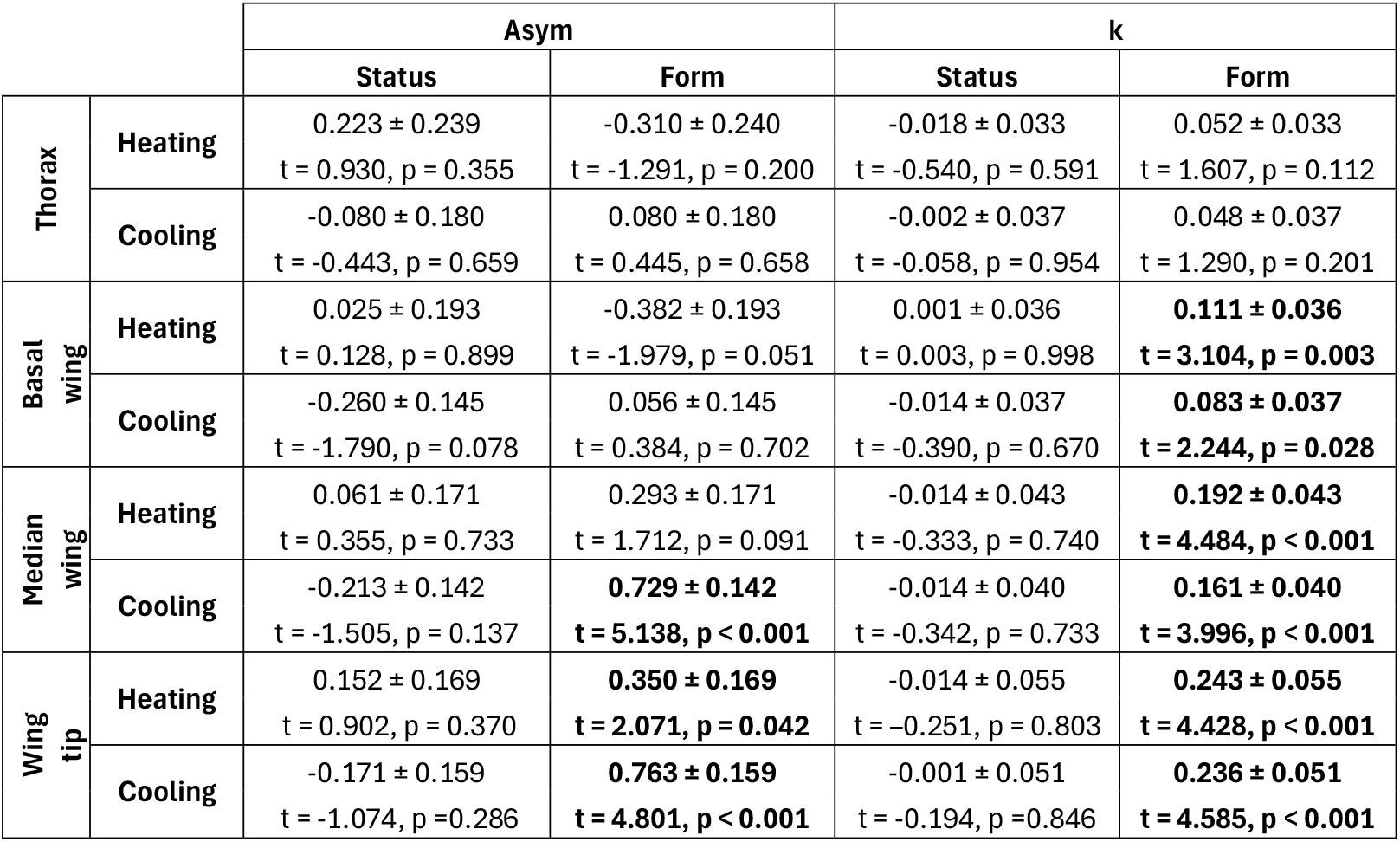
Comparison of model parameters (Asym, k) for thorax and wing tip heating and cooling curves between seasonal forms (spring vs. summer) and specimen status (alive vs. dead) in *Araschnia levana*. Curves were standardised to a common starting temperature and described by asymptotic temperature (Asym) and the curve shape parameter (k), which reflects the steepness of the curve. Linear models were used to test for effects of form and status. Values represent means ± standard error; significant results are marked in **bold**. The reference levels for all linear models were “alive” (for status) and “spring” (for seasonal form), reported mean values represent differences relative to these reference groups (summer – spring, dead – alive).

On the three wing ROIs, however, Asym differed significantly between forms for both wing tip ROI (after heating and after cooling) as well as the median wing ROI after cooling. After heating, wing tip Asym was significantly lower in spring individuals (emmeans contrast: 7.11 ± 0.12°C) compared to summer ones (7.46 ± 0.12°C), conversely summer forms (7.47 ± 0.11°C) had higher Asym after cooling (i.e., more heat loss) compared to spring individuals (6.71 ± 0.11°C), likely related to having gained more heat previously. Wing heating curves of summer individuals rose faster, as indicated by significant differences in k (Table 1). Conversely, cooling curves for summer individuals fell more steeply than for spring ones.

### Morphology drives heating and cooling of the thorax

Higher wing loading had a significant negative effect on heating, i.e. resulted in slower heating (wing loading × time: F = 6.62, p < 0.001, Figure 3A). Seasonal forms differed significantly in time (form × time: F = 2.59, p = 0.037) with the spring form reaching a higher temperature (Table 2). Post-hoc contrasts showed that this interaction was driven by differences between seasonal forms during the heating phase at 60 s (p = 0.037) and 180 s (p = 0.022). The thorax of spring individuals heated ∼0.5 °C more than summer individuals (at 60s and after). Showing that subtle form-related differences emerge only after accounting for wing loading, which strongly influences heating (Figure 3 and 4).

**Table 2:**
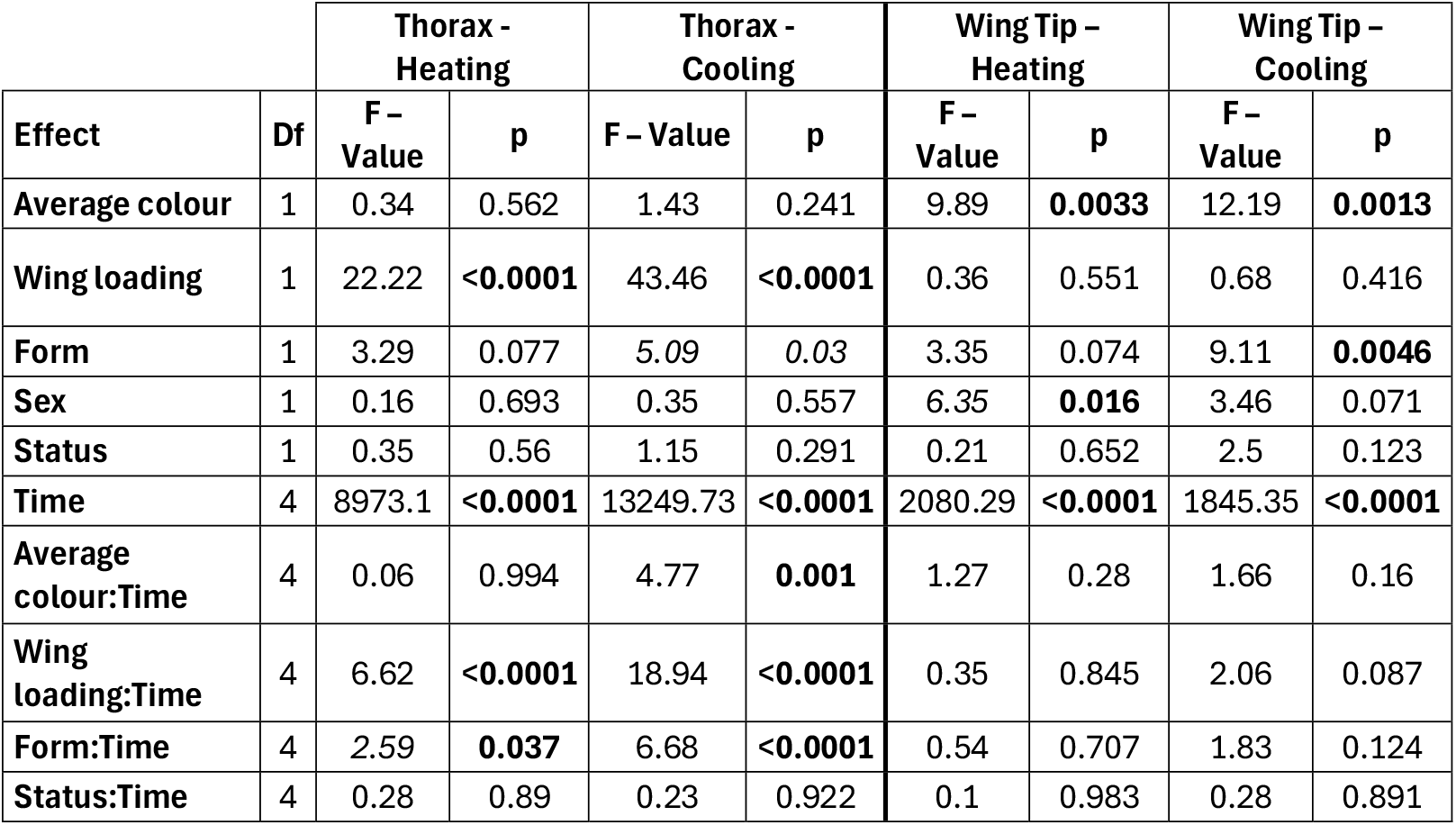
Results of the linear mixed-effect models (2a) for heating and cooling dynamics on the thorax and the wing tip of *Araschnia levana*. Significant effects are marked in **bold**. Output tables for all four ROIs (i.e. thorax and wing basal, median part and wing tip), the two models (wing loading, wing area and weight) as well as contrasts through time for categorical factors are available in Appendix S5.

During cooling, both wing loading and form showed significant effects across time (wing loading × time: F = 18.94, p < 0.001; form × time: F = 6.68, p < 0.001). Consistent with physical principles, higher wing loading resulted in slower cooling through time. Post hoc contrasts indicate that form differences (spring form cooled ∼0.5°C more) emerge only after the initial cooling phase, becoming significant at 60 s (p = 0.012), 180 s (p = 0.002), and 300 s (p = 0.003), while no differences are observed at earlier times (Figure 3). Specimen status (live vs. dead) had no effect. Average colour also showed a significant interaction with time (F = 4.77, p = 0.001). This likely reflects a strong correlation with seasonal form: spring and summer individuals differed in colour, and the form × time effect dominates the dynamics. Complete LMM results can be found in Appendix S5, Tables S3 and S5.

The alternative model using weight and area was consistent with the wing loading model (Appendix S5, Tables S4 and S6). Higher total wing area increased heating and cooling, whereas higher weight had a negative influence.

### Colour drives heating and cooling of the wing tip

For the heating dynamics at the wing tip, none of the parameters (model 2a), showed significant effects over time as heating was primarily driven by average colour (Table 2), with darker individuals heating faster (F = 9.89, p = 0.003). Sex had a significant effect (F = 6.35, p = 0.016) with females reaching higher temperatures compared to males. The effect of form was close to significant (F = 3.35, p = 0.07), driven by a significant difference between spring and summer individuals at 10 s (p = 0.03). At 10 s, wing tips of summer individuals were approx. 0.5 °C warmer (Figure 3).

Cooling of the wing tip showed the same pattern. Colour had a strong effect on cooling, with darker individuals cooling faster (F = 12.19, p = 0.001), as did form (F = 9.11, p = 0.005). Post hoc contrasts indicate that wing tips of spring individuals remained significantly warmer throughout most of the cooling period (10–180s, up to 0.8°C) compared to the summer individuals, with temperature differences decreasing and being similar between forms towards the end (300 s, p = 0.12). Wing loading, sex, and status did not affect cooling thermal dynamics of the wing tip and none of the parameters showed significant effects over time. Model results are in Appendix S5 Tables S7 and S9.

Additional models testing the effect of weight and total wing area model on heating and cooling of the wing tip can be found in Appendix S5 Tables S7 and S9. In both the heating and cooling models, only average colour showed a significant effect. However, the effects across time of average colour, weight and form were close to significance in the cooling model (p = 0.062, 0.089 and 0.051, respectively).

### Heating and cooling of median and basal wing areas is intermediate to the thorax and wing tip

Patterns observed for the basal and median parts of the wing formed a gradient from morphology-dominated in the thorax to colour-dominated at the wing tip.

At the basal wing region (Appendix S5 Table S11-S14), heating rates were generally uniform across forms and colours (also likely due to the basal wing area being black in both forms). In the wing loading model only sex had a significant effect (Appendix S5 Table S11). For the weight and area model, total wing area and the interaction of form and time were significant (Appendix S5 Table S12). The effect of form was driven by differences during the later stages of heating (180 and 300s), with spring individuals reaching on average 0.68°C higher temperature compared to summer ones at the end of heating. Cooling dynamics were significantly correlated with average colour in both models and sex for the wing loading model. Time dependant interactions of form remained insignificant (Appendix S5 Table S13-14).

During heating, the median wing region (Appendix S5 Table S15-S18) showed a significant effect of average colour and form, but their interactions with time were negligible, suggesting parallel heating trajectories across forms. During cooling, both colour and form were highly significant, post-hoc tests indicate summer individuals always cool significantly faster, similar to the effect observed on the wing tip. In both wing regions (median and basal) in cooling models (model 2a or 2b), status (alive vs. dead) had significant effects with wings of live individuals being ∼0.2°C colder than dead ones, however interactions with time remained insignificant. Post-hoc contrasts for the basal wing were significant during later stages of cooling (180 and 300s) but remained insignificant for the median wing through all-time points except in the weight and area model at 300s (Figure 3).

## Discussion

In *Araschnia levana*, the thoracic cooling and heating dynamics are primarily shaped by wing loading differences between individuals (Table 2), while thermal dynamics of the wings were determined by individual wing coloration (Table 1 & 2). The interaction between form-specific morphology in wing loading and colouration resulted in overall similar thoracic thermal dynamics between the seasonal forms (Table 1, Figure 2). After accounting for the effects of wing loading and colour, we identified differences in heating and cooling dynamics between the two seasonal forms (Figure 3 & 4). This likely reflects unmeasured physiological or anatomical differences related to climatic seasonality (Baudach et al., 2018). The polyphenism of *A. levana* is therefore unlikely to be directly driven by thermoregulation demands but seasonal forms show optimised thermal performance.

**Figure 2:**
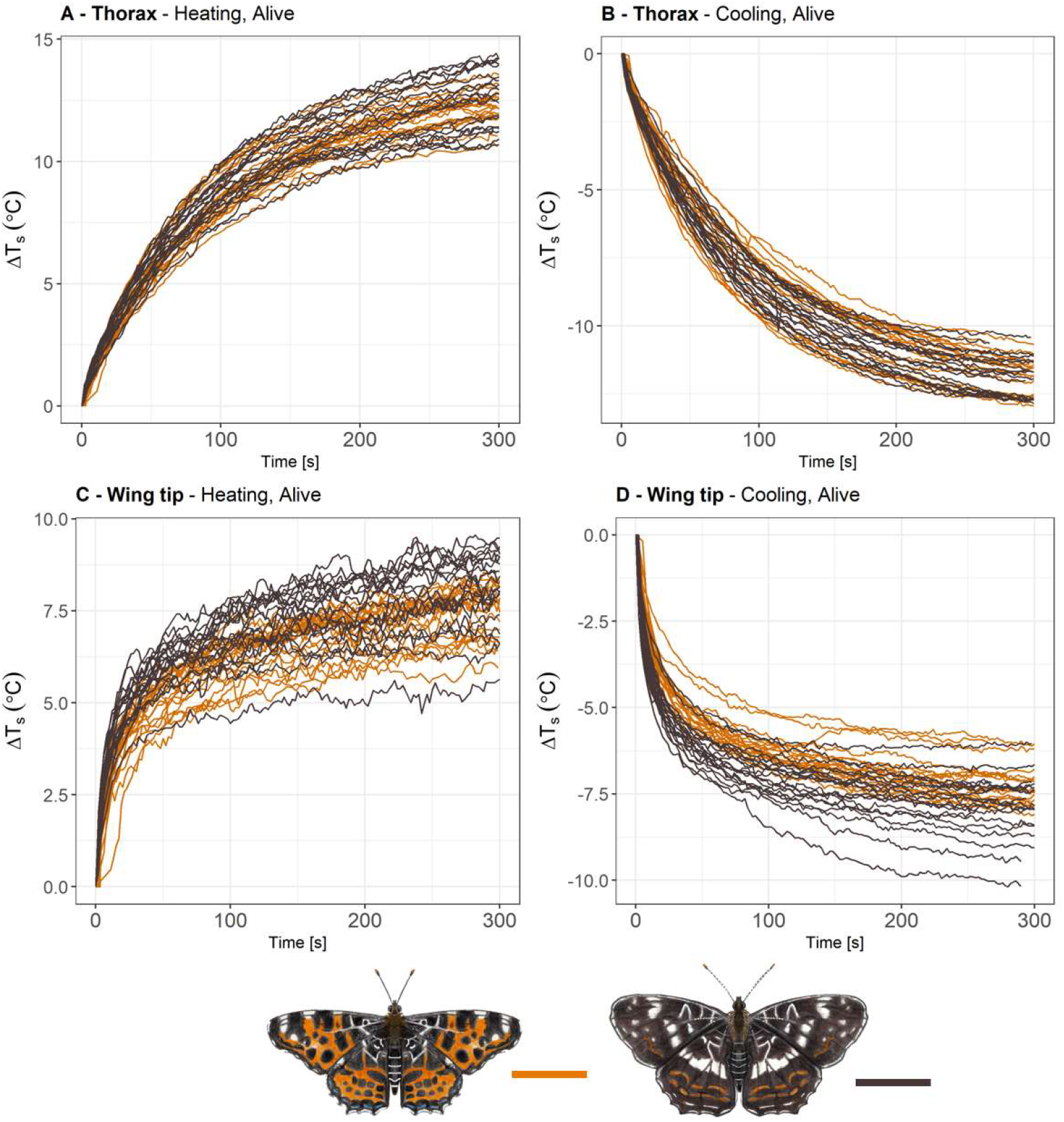
Thoracic heating and cooling of (**A, B**) in *Araschnia levana* did not differ between spring and summer form, nor between alive and dead butterflies. (**C**) On wing tips, curves of summer individuals reached higher asymptotic temperatures (Asym) and rose faster, while cooling curves (**D**) reached lower Asym and fell faster compared to the spring form. Thorax and wing temperatures were standardised using ΔT compared to T_0_, separately for heating and cooling. Plots for all ROIs and curves comparing dead and alive individuals can be found in Appendix S4, Figure S1-4).

### Morphological differences between forms

We confirmed significant morphometric differences between the seasonal forms, consistent with earlier studies (Fric et al., 2006; Esperk & Tammaru, 2021). Summer individuals exhibited significantly larger wings and lower wing loading than spring forms. The seasonal difference in wing loading aligns with patterns seen in other Lepidoptera where summer morphs have longer wings (Komata & Sota, 2017; Kleckova et al., 2024) and are better suited for dispersal or rapid movements in warm, resource rich environments (Fric & Konvicka, 2002; Sekar, 2012). Conversely, the spring form’s higher wing loading and reduced wing area may reflect trade-offs favouring thermal efficiency and resource conservation in cooler, more constrained early-season environments.

### Absence of active thermoregulation by haemolymph circulation

By comparing live and dead individuals, we assessed whether haemolymph circulation affects heating and cooling in *A. levana* in our experimental setup. We found no differences in thoracic heating and cooling rates between live and dead individuals. Similarly, Wasserthal (1982) with a comparable setup did not reveal differences in thermal dynamics between dead and alive pierid butterflies. We however observed significant differences during cooling on the median wing between live and dead specimens, where live specimens were roughly 0.2°C colder than dead ones (Appendix S5 Table S13, S14 and S17). However, as we did not record a significant difference in thorax temperature, which is the biologically relevant proxy of butterfly activity, we do not expect a major effect of haemolymph circulation on thermoregulation in *A. levana*. The role of active haemolymph circulation by compressing the tracheal system (for review: Salcedo & Socha, 2020) has been confirmed to contribute to thermoregulation in the Atlas moth (Wasserthal, 1982) and during overheating in small/medium sized butterflies where active cooling of the wings was detected (Tsai et al., 2020). In our experimental setup, we suppressed the subtle behavioural mechanisms affecting thermoregulation, such as wing positioning, or wing movement (Kingsolver, 1995; Dongmo et al., 2018). Future work should test these thermal dynamics under field-realistic conditions, integrating behavioural thermoregulation (e.g., basking posture, timing of daily activity).

### Colour and wing loading are main drivers of thermal dynamics

Wing loading and colouration were the main drivers of heating and cooling dynamics on the thorax and along the wing of *A. levana*. Higher wing loading slows down heating by reducing heat loss. Similarly, Kleckova, et al. (2023b) identified wing loading as the main determinant of heating in *Erebia* mountain butterflies. Generally, heavier individuals heated and cooled slower, consistent with the thermal inertia conferred by greater weight. In contrast, larger wing area accelerated both heating and cooling. On the wing, darker individuals heated up faster initially as expected based on basic physical principles, this however did not affect thoracic temperature. Similarly, in a lycaenid butterfly, *Polyommatus icarus*, body size rather than wing melanisation determined heating rates (De Keyser et al., 2015). However, darker wing coloration of butterflies was linked to more effective thorax heating in a community of tropical butterflies (Ashe-Jepson et al., 2023). While thermoregulation may represent a minor or indirect selective pressure underlying polyphenism in *A. levana*, it does not appear to be its primary driver.

### Thermal dynamics of seasonal forms differ in individuals with similar morphology

When morphology is considered, significant differences in thermal dynamics emerge between spring and summer forms, both on the thorax and on the wing (Figures 3 & 4). Seasonal forms thus differ in their heating and cooling dynamics in ways not explained by morphology and colouration alone. We show that the thorax of orange spring individuals heated and cooled by ∼0.5°C more than that of the black summer ones. This suggests that thermal dynamics might be affected by factors beyond the scope of our study. For example, differences in cellular structure such as membrane lipid composition (Slotsbo et al., 2016; Rodríguez et al., 2018) or cuticular structure characterised by its thickness or microstructure (López et al., 2024; Yao et al., 2025) as observed in beetles (Amore et al., 2017; Pavlović et al., 2018) might affect thermoregulation. Alternatively, heterogenous variation in scale arrangement resulting in varying emissivity, was shown to improve cooling in a tropical butterfly, *Tirumala limniace* (Tang et al., 2022). Colour patterns might also affect observed thermal differences between forms in individuals of standard size. For example, the white disruptive bands in the subtropical butterfly *Anartia fatima*, similar as in the summer form of *A. levana*, reduces heating and counters heat stress (Brashears et al., 2016), thus potentially promoting the reduction of heating rates observed in summer individuals of *A. levana*.

**Figure 3:**
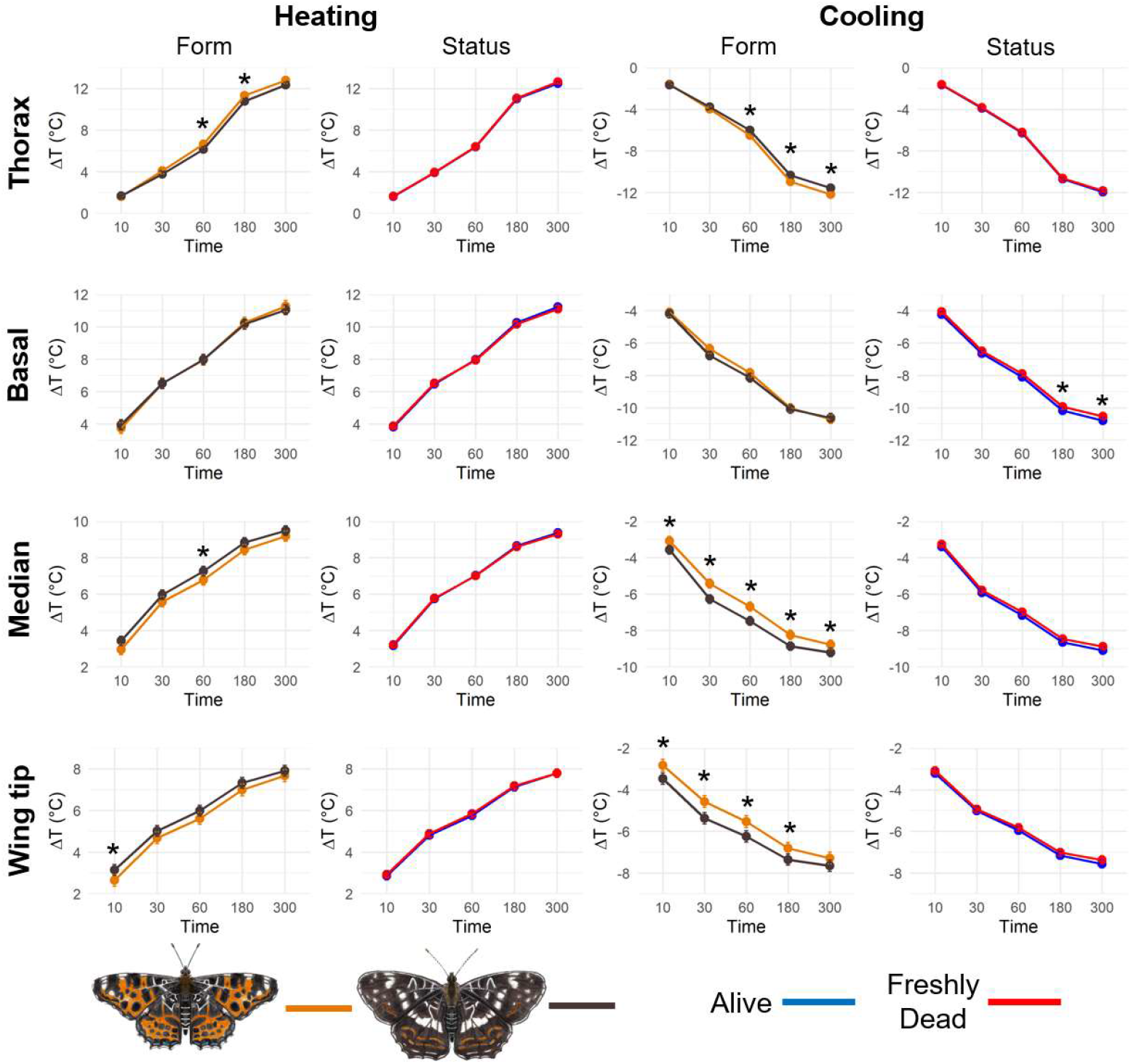
Model-adjusted temperature means (model 2a) through time for temperature dynamics of *Araschnia levana* across all four tested ROIs, for seasonal form (spring vs. summer) and specimen status (live vs. dead). Temperature differences (ΔT) represent the change in each ROI at multiple time points (10, 30, 60, 180, and 300 seconds since experiment start) relative to the baseline temperature, T_0_. Asterisks (*) indicate significant differences between classes based on emmeans pairwise contrasts. Note that the x-axis is not linear.

**Figure 4:**
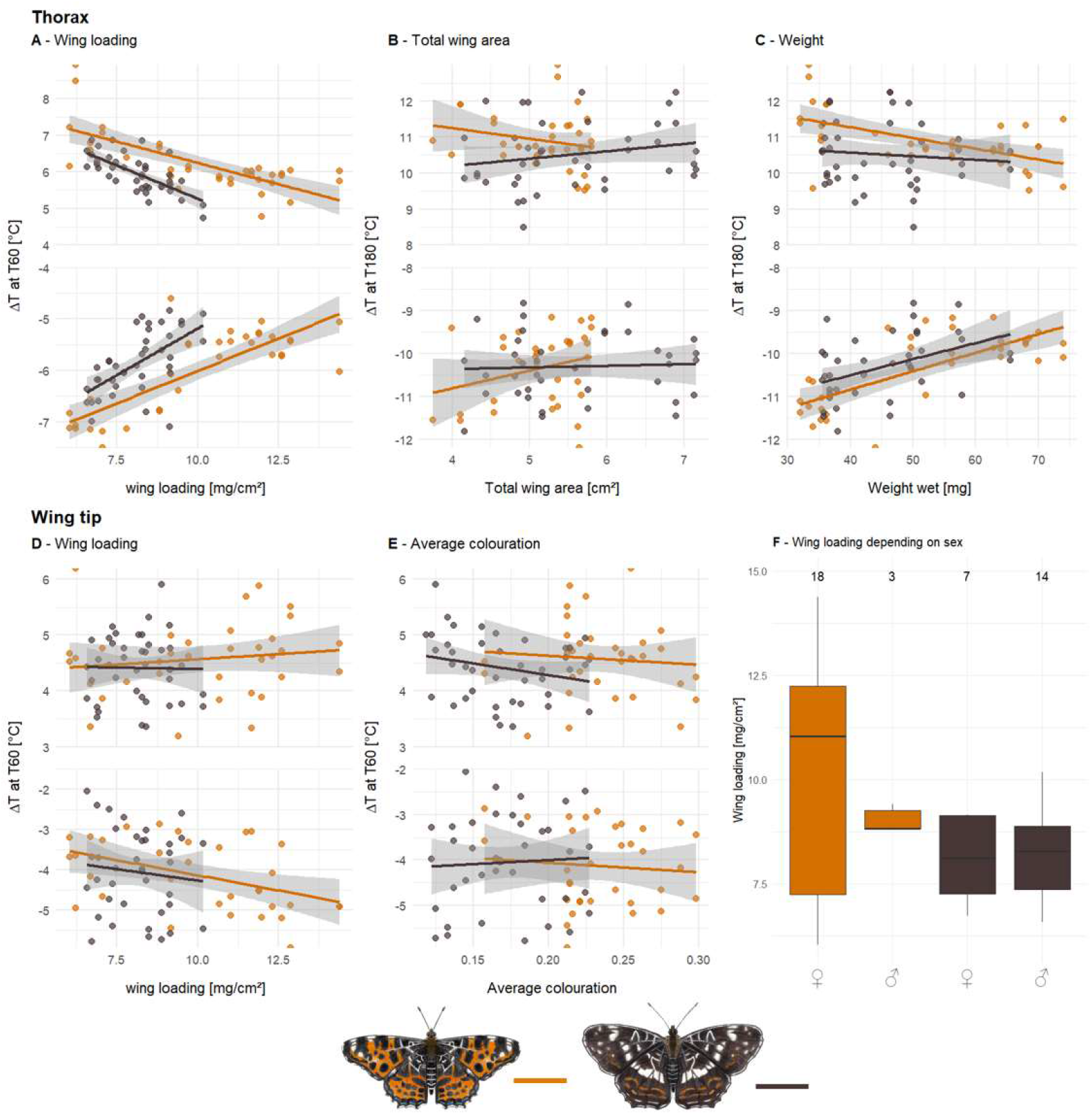
Data plots showing the relationship between heating and cooling dynamics and morphometric parameters in the spring and summer forms of *Araschnia levana* (**A-E**). The upper parts of the panels illustrate heating dynamics (positive ΔT differences), while the lower parts show cooling dynamics (negative ΔT differences). **A-C** depict thermal dynamics on the thorax; **D-E** thermal dynamics on the wing tip. **F** shows how wing loading differs between forms depending on sex. Data combines dead and alive specimens as no differences between them were found (i.e., two observations per specimen).

The observed differences between forms in individuals of standard size might relate to seasonal conditions. Spring individuals are expected to be under selection to heat up faster and remain active under colder and more unpredictable spring conditions in agreement with more effective heating observed in spring individuals of *A. levana* (Figure 4). The faster heat loss during cooling in spring individuals may simply be an indirect consequence of ability to heat up more effectively. This heat loss might be balanced by higher wing loading of the spring form, which enables heat accumulation in the body and lower loss due to smaller wing area. In contrast, summer individuals are in flight during warmer conditions. Thus, their thermal dynamics are slower, despite lower wing loading, reflecting that rapid heating or cooling is not critical during summer. As heating start conditions were standardised separately for heating and cooling phases, cooling asymptotes partly reflect heat gained during the heating phase, meaning that differences during cooling may originate from differences during heating. This, however, does not affect interpretation of overall form-specific thermal dynamics.

It is expected for summer individuals to heat up faster, due to their dark colouration (Clusella-Trullas et al., 2008; Xing et al., 2016; Järvi et al., 2019; Bladon et al., 2020), however this was only significant during the first time point on the wing tip at 10s of heating, likely due to summer individuals absorbing more heat initially (Figure 3), but this effect dissipated as heating progressed. This indicates that despite being significantly lighter in colour, wings of spring individuals can achieve similar heating rates as darker wings in the summer form (Figure 4). Surprisingly, the wings of summer individuals cooled significantly faster at almost all time points (excluding 300 s), which may reflect an adaptation to summer climate, allowing the wings to emit heat more efficiently and avoid overheating (despite the higher heat absorption caused by dark colouration). Potential causes might be the presence of a disruptive cooling band, scale arrangements or cuticular differences as discussed earlier (Brashears et al., 2016; Amore et al., 2017; Pavlović et al., 2018; Tang et al., 2022).

We expect that the seasonal polyphenism of *A. levana* is not adaptive in relation to thermal dynamics. The summer phenotype of *A. levana* may represent an ancestral mimetic trait driven by predation as suggested by Fric et al. (2004). They hypothesised that the darker colouration of the summer form evolved in seasonal tropical regions, potentially involving mimicry of Limenitidae species (e.g., *Neptis* spp.), which are highly speciose in Southeast Asia and exhibit similar wing patterns. It would be interesting to test whether related species in the genus *Araschnia*, occurring in tropical regions, or other multivoltine temperate species, show similar thermal dynamics.

## Conclusions

Our findings demonstrate that *A. levana’s* seasonal size and colour polyphenism while affecting thermal performance is unlikely to be the driver of the present polyphenism. It is already counterintuitive for the black form to fly during the warmer season as more melanistic colouration is usually an adaptation observed in colder conditions, i.e., higher altitudes/latitudes or colder seasons. When accounting for differences in wing loading, the black summer form heats up and cools down slower than the lighter orange form. This contra-intuitive pattern suggests for seasonal optimisation of thermoregulation. Further mechanistic investigations of cuticle thickness, surface microstructure, scale heterogeneity and radiation reflectance might explain the unexpected connections between morphology, colouration, and thermoregulation.

## Acknowledgment

We would like to thank Michaela Helclová who created the butterfly and experimental setup drawings used throughout the manuscript.

## Authors contribution

**DL** - study planning, field investigation, experiment, data evaluation, writing first draft, review and editing; **JO** - study planning, data evaluation, review and editing; **AS** - field investigation, review and editing; **LJ** – data evaluation, review and editing; **PM** - review and editing, funding acquisition; **IK** - study planning, field investigation, experiment, review and editing, funding acquisition.

## Open research statement

The data that support the findings of this study are available from the corresponding author upon request and will be archived on Zenodo after acceptance.

## References

Amore, V., Hernández, M. I. M., Carrascal, L. M., & Lobo, J. M. (2017). Exoskeleton may influence the internal body temperatures of Neotropical dung beetles (Col. Scarabaeinae). PeerJ, 5, e3349. 10.7717/peerj.3349

Ashe-Jepson, E., Arizala Cobo, S., Basset, Y., Bladon, A. J., Kleckova, I., Laird-Hopkins, B. C., Mcfarlane, A., Sam, K., Savage, A. F., Zamora, A. C., Turner, E. C., & Lamarre, G. P. A. (2023). Tropical butterflies use thermal buffering and thermal tolerance as alternative strategies to cope with temperature increase. Journal of Animal Ecology, 92(9), 1759–1770. 10.1111/1365-2656.13970

Bates, D., Mächler, M., Bolker, B., & Walker, S. (2015). Fitting Linear Mixed-Effects Models Using lme4. Journal of Statistical Software, 67, 1–48. 10.18637/jss.v067.i01

Baudach, A., Lee, K.-Z., Vogel, H., & Vilcinskas, A. (2018). Immunological larval polyphenism in the map butterfly Araschnia levana reveals the photoperiodic modulation of immunity. Ecology and Evolution, 8(10), 4891–4898. 10.1002/ece3.4047

Bladon, A. J., Lewis, M., Bladon, E. K., Buckton, S. J., Corbett, S., Ewing, S. R., Hayes, M. P., Hitchcock, G. E., Knock, R., Lucas, C., McVeigh, A., Menéndez, R., Walker, J. M., Fayle, T. M., & Turner, E. C. (2020). How butterflies keep their cool: Physical and ecological traits influence thermoregulatory ability and population trends. Journal of Animal Ecology, 89(11), 2440–2450. 10.1111/1365-2656.13319

Brashears, J., Aiello, A., & Seymoure, B. M. (2016). Cool Bands: Wing bands decrease rate of heating, but not equilibrium temperature in Anartia fatima. Journal of Thermal Biology, 56, 100–108. 10.1016/j.jtherbio.2016.01.007

Cleveland, W. S. (1979). Robust Locally Weighted Regression and Smoothing Scatterplots. Journal of the American Statistical Association, 74(368), 829–836. 10.1080/01621459.1979.10481038

Clusella-Trullas, S., Terblanche, J. S., Blackburn, T. M., & Chown, S. L. (2008). Testing the thermal melanism hypothesis: A macrophysiological approach. Functional Ecology, 22(2), 232–238. 10.1111/j.1365-2435.2007.01377.x

De Keyser, R., Breuker, C. J., Hails, R. S., Dennis, R. L. H., & Shreeve, T. G. (2015). Why Small Is Beautiful: Wing Colour Is Free from Thermoregulatory Constraint in the Small Lycaenid Butterfly, Polyommatus icarus. PLoS ONE, 10(4), e0122623. 10.1371/journal.pone.0122623

Dongmo, M. A. K., Bonebrake, T. C., Hanna, R., & Fomena, A. (2018). Seasonal Polyphenism in Bicyclus dorothea (Lepidoptera: Nymphalidae) Across Different Habitats in Cameroon. Environmental Entomology, 47(6), 1601–1608. 10.1093/ee/nvy135

Esperk, T., & Tammaru, T. (2021). Ontogenetic Basis of Among-Generation Differences in Size-Related Traits in a Polyphenic Butterfly. Frontiers in Ecology and Evolution, 9. 10.3389/fevo.2021.612330

Fric, Z., Klimova, M., & Konvicka, M. (2006). Mechanical design indicates differences in mobility among butterfly generations. Evolutionary Ecology Research, 8(8), 1511–1522.

Fric, Z., & Konvicka, M. (2002). Generations of the polyphenic butterfly Araschnia levana differ in body design. Evolutionary Ecology Research, 4(7), 1017–1032.

Fric, Z., Konvicka, M., & Zrzavy, J. (2004). Red & black or black & white? Phylogeny of the Araschnia butterflies (Lepidoptera: Nymphalidae) and evolution of seasonal polyphenism. Journal of Evolutionary Biology, 17(2), 265–278.

Halali, S., Brakefield, P. M., & Brattström, O. (2024). Phenotypic plasticity in tropical butterflies is linked to climatic seasonality on a macroevolutionary scale. Evolution, 78(7), 1302–1316. 10.1093/evolut/qpae059

Järvi, V. V., Burg, K. R. L. van der, & Reed, R. D. (2019). Seasonal Plasticity in Junonia coenia (Nymphalidae): Linking Wing Color, Temperature Dynamics, and Behavior. The Journal of the Lepidopterists’ Society, 73(1), 34–42. 10.18473/lepi.73i1.a5

Kemp, D. J., & Krockenberger, A. K. (2002). A novel method of behavioural thermoregulation in butterflies. Journal of Evolutionary Biology, 15(6), 922–929. 10.1046/j.1420-9101.2002.00470.x

Kingsolver, J. G. (1995). Fitness consequences of seasonal polyphenism in western white butterflies. Evolution, 49(5), 942–954.

Kleckova, I., Klečka, J., Fric, Z., česanek, M., Dutoit, L., Pellissier, L., & Matos-Maraví, P. (2023a). Climatic Niche Conservatism and Ecological Diversification in the Holarctic Cold-Dwelling Butterfly Genus Erebia. Insect Systematics and Diversity, 7. 10.1093/isd/ixad002

Kleckova, I., Konvicka, M., & Klecka, J. (2014). Thermoregulation and microhabitat use in mountain butterflies of the genus Erebia: Importance of fine-scale habitat heterogeneity. Journal of Thermal Biology, 41, 50–58. 10.1016/j.jtherbio.2014.02.002

Kleckova, I., Linke, D., Rezende, F. D. M., Rauscher, L., Le Roy, C., & Matos-Maraví, P. (2024). Flight behaviour diverges more between seasonal forms than between species in Pieris butterflies. Ecology and Evolution, 14(7), e70012. 10.1002/ece3.70012

Kleckova, I., Okrouhlík, J., Svozil, T., Matos-Maraví, P., & Klecka, J. (2023b). Body size, not species identity, drives body heating in alpine Erebia butterflies. Journal of Thermal Biology, 113, 103502. 10.1016/j.jtherbio.2023.103502

Komata, S., & Sota, T. (2017). Seasonal polyphenism in body size and juvenile development of the swallowtail butterfly Papilio xuthus (Lepidoptera: Papilionidae). EJE, 114(1), 365–371. 10.14411/eje.2017.046

Lahondère, C. (2023). Recent advances in insect thermoregulation. Journal of Experimental Biology, 226(18), jeb245751. 10.1242/jeb.245751

Laird-Hopkins, B. C., Ashe-Jepson, E., Basset, Y., Arizala Cobo, S., Eberhardt, L., Freiberga, I., Hellon, J., Hitchcock, G. E., Kleckova, I., Linke, D., Lamarre, G. P. A., McFarlane, A., Savage, A. F., Turner, E. C., Zamora, A. C., Sam, K., & Bladon, A. J. (2023). Thermoregulatory ability and mechanism do not differ consistently between neotropical and temperate butterflies. Global Change Biology, 29(15), 4180–4192. 10.1111/gcb.16797

Lenth, R. (2025). emmeans: Estimated Marginal Means, aka Least-Squares Means. (Version 1.11.1-00001) [Computer software]. https://rvlenth.github.io/emmeans/

Linke, D., Debat, V., & Matos-Maravi, P. (2025). Association of iridescence and tails with flight proxies suggests that predation drives convergence in skipper butterflies (p. 2025.06.23.661062). bioRxiv. 10.1101/2025.06.23.661062

López, Q. K., Cárdenas, R. E., Castro, F. R., Vizuete, K., Checa, M. F., Vera, C. C., López, Q. K., Cárdenas, R. E., Castro, F. R., Vizuete, K., Checa, M. F., & Vera, C. C. (2024). Nanostructural Influence on Optical and Thermal Properties of Butterfly Wing Scales Across Forest Vertical Strata. Materials, 17(20). 10.3390/ma17205084

Markl, G., Ottmann, S., Haasis, T., Budach, D., Krais, S., & Köhler, H.-R. (2022). Thermobiological effects of temperature-induced color variations in Aglais urticae (Lepidoptera, Nymphalidae). Ecology and Evolution, 12(6), e8992. 10.1002/ece3.8992

Nelson, R. J., Denlinger, D. L., & Somers, D. E. (2010). Photoperiodism: The biological calendar. Oxford University Press.

Pavlović, D., Vasiljević, D., Salatić, B., Lazović, V., Dikić, G., Tomić, L., Ćurčić, S., Milovanović, P., Todorović, D., & Pantelić, D. V. (2018). Photonic structures improve radiative heat exchange of Rosalia alpina (Coleoptera: Cerambycidae). Journal of Thermal Biology, 76, 126–138. 10.1016/j.jtherbio.2018.07.014

Pinheiro, J., Bates, D., & R Core Team. (2025). nlme: Linear and Nonlinear Mixed Effects Models. https://CRAN.R-project.org/package=nlme

R Core Team. (2025). R: A Language and Environment for Statistical Computing. R Foundation for Statistical Computing. https://www.R-project.org/

Rodríguez, E., Weber, J.-M., & Darveau, C.-A. (2018). Diversity in membrane composition is associated with variation in thermoregulatory capacity in hymenopterans. Comparative Biochemistry and Physiology. Part B, Biochemistry & Molecular Biology, 224, 115–120. 10.1016/j.cbpb.2017.11.017

Rosa, E., & Saastamoinen, M. (2020). Beyond thermal melanism: Association of wing melanization with fitness and flight behaviour in a butterfly. Animal Behaviour, 167, 275–288. 10.1016/j.anbehav.2020.07.015

Salcedo, M. K., & Socha, J. J. (2020a). Circulation in Insect Wings. Integrative and Comparative Biology, 60(5), 1208–1220. 10.1093/icb/icaa124

Salcedo, M. K., & Socha, J. J. (2020b). Circulation in Insect Wings. Integrative and Comparative Biology, 60(5), 1208–1220. 10.1093/icb/icaa124

Schneider, C. A., Rasband, W. S., & Eliceiri, K. W. (2012). NIH Image to ImageJ: 25 years of image analysis. Nature Methods, 9(7), Article 7. 10.1038/nmeth.2089

Sekar, S. (2012). A meta-analysis of the traits affecting dispersal ability in butterflies: Can wingspan be used as a proxy? Journal of Animal Ecology, 81(1), 174–184. 10.1111/j.1365-2656.2011.01909.x

Slotsbo, S., Sørensen, J. G., Holmstrup, M., Kostal, V., Kellermann, V., & Overgaard, J. (2016). Tropical to subpolar gradient in phospholipid composition suggests adaptive tuning of biological membrane function in drosophilids. Functional Ecology, 30(5), 759–768. 10.1111/1365-2435.12568

Tang, C.-F., Li, F.-F., Cao, Y., & Liao, H.-J. (2022). Universal cooling patterns of the butterfly wing scales hierarchy deduced from the heterogeneous thermal and structural properties of Tirumala limniace (Lepidoptera: Nymphalidae, Danainae). Insect Science, 29(6), 1761–1772. 10.1111/1744-7917.13046

Tsai, C.-C., Childers, R. A., Nan Shi, N., Ren, C., Pelaez, J. N., Bernard, G. D., Pierce, N. E., & Yu, N. (2020). Physical and behavioral adaptations to prevent overheating of the living wings of butterflies. Nature Communications, 11(1), 551. 10.1038/s41467-020-14408-8

Van Dyck, H., & Wiklund, C. (2002). Seasonal butterfly design: Morphological plasticity among three developmental pathways relative to sex, flight and thermoregulation. Journal of Evolutionary Biology, 15(2), 216–225. 10.1046/j.1420-9101.2002.00384.x

Vilcinskas, A., & Vogel, H. (2016). Seasonal phenotype-specific transcriptional reprogramming during metamorphosis in the European map butterfly Araschnia levana. Ecology and Evolution, 6(11), 3476–3485. 10.1002/ece3.2120

Wang, L.-Y., Franklin, A. M., Black, J. R., & Stuart-Fox, D. (2021). Heating rates are more strongly influenced by near-infrared than visible reflectance in beetles. Journal of Experimental Biology, 224(19), jeb242898.

Wasserthal, L. T. (1982). Antagonism between haemolymph transport and tracheal ventilation in an insect wing (Attacus atlas L.). Journal of Comparative Physiology, 147(1), 27–40. 10.1007/BF00689287

Xing, S., Bonebrake, T. C., Tang, C. C., Pickett, E. J., Cheng, W., Greenspan, S. E., Williams, S. E., & Scheffers, B. R. (2016). Cool habitats support darker and bigger butterflies in Australian tropical forests. Ecology and Evolution, 6(22), 8062–8074. 10.1002/ece3.2464

Yao, K., Kong, G., Xiao, C., Chen, S., Zhang, Y., Lou, X., Li, J., Zhang, D., Zhou, H., & Zheng, Y. (2025). Bioinspired photonic materials for advanced thermal management. Chemical Society Reviews, 54(22), 10690–10723. 10.1039/D5CS00471C

