## Appendix for "Heated debate: Is the seasonal polyphenism of *Araschnia levana* (Lepidoptera: Nymphalidae) driven by thermoregulation?"

| | |
|---|---|
| Daniel Linke: | 0000-0002-0686-3961 |
| Jan Okrouhlik: | 0000-0001-9935-7859 |
| Alena Sucháčková Bartoňová: | 0000-0001-6298-2466 |
| Leonardo Ré Jorge: | 0000-0003-4518-4328 |
| Pável Matos-Maraví: | 0000-0002-2885-4919 |
| Irena Klečková: | 0000-0002-9333-213X |

**Appendix S1: Sampling localities**

**Table S1:** Full list of utilised specimens indicating collection date, locality, sex and form.

| ID | Collection date | Locality | Coordinates | Sex | Form |
|---|---|---|---|---|---|
| L001 | 20.05.2023 | Český Krumlov | 48.82170, 14.27971 | male | spring |
| L002 | 21.05.2023 | Halamky | 48.85097, 14.91245 | male | spring |
| L003 | 21.05.2023 | Halamky | 48.85097, 14.91245 | female | spring |
| L004 | 25.05.2023 | České Budějovice | 48.96835, 14.43369 | female | spring |
| L005 | 25.05.2023 | České Budějovice | 48.96835, 14.43369 | female | spring |
| L006 | 28.05.2023 | Český Krumlov | 48.82170, 14.27971 | female | spring |
| L007 | 07.06.2023 | Mariánské Lázně | 49.98838, 12.72124 | female | spring |
| L008 | 09.06.2023 | Mariánské Lázně | 49.98838, 12.72124 | female | spring |
| L009 | 09.06.2023 | Mariánské Lázně | 49.98838, 12.72124 | female | spring |
| L010 | 09.06.2023 | Mariánské Lázně | 49.98838, 12.72124 | female | spring |
| L011 | 09.06.2023 | Mariánské Lázně | 49.98838, 12.72124 | female | spring |
| L012 | 09.06.2023 | Mariánské Lázně | 49.98838, 12.72124 | female | spring |
| L013 | 09.06.2023 | Mariánské Lázně | 49.98838, 12.72124 | female | spring |
| L014 | 09.06.2023 | Mariánské Lázně | 49.98838, 12.72124 | female | spring |
| L015 | 09.06.2023 | Mariánské Lázně | 49.98838, 12.72124 | female | spring |
| L016 | 09.06.2023 | Mariánské Lázně | 49.98838, 12.72124 | female | spring |
| L017 | 09.06.2023 | Mariánské Lázně | 49.98838, 12.72124 | female | spring |
| L018 | 09.06.2023 | Mariánské Lázně | 49.98838, 12.72124 | female | spring |
| L019 | 09.06.2023 | Mariánské Lázně | 49.98838, 12.72124 | female | spring |
| L020 | 09.06.2023 | Mariánské Lázně | 49.98838, 12.72124 | male | spring |
| L021 | 09.06.2023 | Mariánské Lázně | 49.98838, 12.72124 | female | spring |
| L047 | 26.07.2023 | České Budějovice | 48.96835, 14.43369 | female | summer |
| L048 | 26.07.2023 | České Budějovice | 48.96835, 14.43369 | male | summer |
| L049 | 26.07.2023 | České Budějovice | 48.96835, 14.43369 | female | summer |
| L058 | 27.07.2023 | České Budějovice | 48.96835, 14.43369 | female | summer |
| L059 | 27.07.2023 | České Budějovice | 48.96835, 14.43369 | female | summer |
| L060 | 27.07.2023 | České Budějovice | 48.96835, 14.43369 | male | summer |
| L066 | 31.08.2023 | Třeboň | 48.92600, 14.77739 | male | summer |
| L067 | 31.08.2023 | Třeboň | 48.92600, 14.77739 | female | summer |
| L068 | 31.08.2023 | Třeboň | 48.92600, 14.77739 | female | summer |
| L069 | 31.08.2023 | Třeboň | 48.92600, 14.77739 | male | summer |
| L070 | 31.08.2023 | Třeboň | 48.92600, 14.77739 | female | summer |
| L071 | 31.08.2023 | Třeboň | 48.92600, 14.77739 | male | summer |
| L072 | 10.08.2023 | Horní Stropnice | 48.74967, 14.71133 | male | summer |
| L073 | 10.08.2023 | Horní Stropnice | 48.74967, 14.71133 | male | summer |
| L074 | 10.08.2023 | Horní Stropnice | 48.74967, 14.71133 | male | summer |
| L075 | 10.08.2023 | Horní Stropnice | 48.74967, 14.71133 | male | summer |

| | | | | | |
|---|---|---|---|---|---|
| L076 | 10.08.2023 | Horní Stropnice | 48.74967, 14.71133 | male | summer |
| L077 | 10.08.2023 | Horní Stropnice | 48.74967, 14.71133 | male | summer |
| L078 | 10.08.2023 | Horní Stropnice | 48.74967, 14.71133 | male | summer |
| L079 | 10.08.2023 | Horní Stropnice | 48.74967, 14.71133 | male | summer |
| L080 | 10.08.2023 | Horní Stropnice | 48.74967, 14.71133 | male | summer |

**Appendix S2:** Thermal data quality control

All thermographic analyses were conducted using a fixed emissivity value of 0.95, consistent with previous studies on Lepidoptera wings and cuticle (Tsai et al., 2020). Emissivity was not independently validated; therefore, absolute temperature estimates may include a small systematic offset. However, because all measurements were obtained under identical conditions, comparisons between body regions and experimental groups remain robust.

To ensure that recorded temperatures reflected emitted surface radiation rather than reflected lamp radiation, we inspected the initial frames immediately following lamp activation. No instantaneous step changes exceeding measurement noise was detected in any ROI, indicating a negligible contribution of reflected radiation within the long-wave infrared band captured by the camera.

To monitor potential systematic bias or temporal drift in the thermal camera, a fine wire thermocouple was placed within the camera's field of view and covered with a thin layer of filter paper to provide a stable, high-emissivity reference surface. Although this reference allowed us to verify measurement stability across trials, it was not used to correct camera readings. All analyses were based on temperature change relative to the pre-heating baseline ($\Delta T$), which effectively cancels any constant offset associated with camera calibration.

Trials were excluded from further analysis if the butterfly exhibited visible movement during recording, if experimental positioning was disrupted, or if ambient room temperature exceeded the predefined threshold of 12.5°C.

**Appendix S3:** Comparison of morphometric measurements between forms and sexes of *A. levana*

**Table S2:** Comparison between forewing length, total wing area, wing loading and average wing colour depending on sex (male vs. female) and form (spring vs. summer), comparisons were done using a linear model and emmeans contrasts. Samples sizes were not equal between sexes. Significant effects marked in **bold**.

| Trait | Factor | Group Means ± SD | LM t (df=1,39) | p-value | Emmeans diff | 95% CI (lwr–upr) |
|---|---|---|---|---|---|---|
| **Forewing length [cm]** | **Form** | Spring: 1.85 ± 0.10<br>Summer: 2.06 ± 0.12 | 6.71 | **<0.001** | 0.215 (Summer > Spring) | Spring: 1.80 – 1.89<br>Summer: 2.02 – 2.10 |
| | **Sex** | Female: 2.05 ± 0.12<br>Male: 1.86 ± 0.11 | -5.72 | **<0.001** | −0.187 (Male < Female) | Female: 2.01 – 2.09<br>Male: 1.81 – 1.91 |
| **Weight [mg]** | **Form** | Spring: 45.3 ± 10.3<br>Summer: 47.5 ± 10.2 | 0.56 | 0.576 | 2.12 (Summer > Spring) | Spring: 40.0 – 50.7<br>Summer: 42.7 – 52.2 |
| | **Sex** | Female: 53.5 ± 10.1<br>Male: 39.3 ± 10.8 | -3.69 | **0.001** | −14.12 (Male < Female) | Female: 49.0 – 58.0<br>Male: 33.7 – 45.0 |
| **Total wing area [cm$^2$]** | **Form** | Spring: 4.58 ± 0.40<br>Summer: 5.80 ± 0.45 | 6.74 | **<0.001** | 1.22 (Summer > Spring) | Spring: 4.32 – 4.84<br>Summer: 5.57 – 6.03 |
| | **Sex** | Female: 5.92 ± 0.44<br>Male: 4.46 ± 0.50 | -7.90 | **<0.001** | −1.46 (Male < Female) | Female: 5.70 – 6.14<br>Male: 4.19 – 4.73 |
| **Wing loading [mg cm$^{-2}$]** | **Form** | Spring: 9.74 ± 1.35<br>Summer: 8.21 ± 1.28 | -2.22 | **0.032** | −1.53 (Summer < Spring) | Spring: 8.76 – 10.72<br>Summer: 7.34 – 9.07 |
| | **Sex** | Female: 9.16 ± 1.28<br>Male: 8.79 ± 1.41 | -0.53 | 0.601 | −0.37 (Male < Female) | Female: 8.33 – 9.98<br>Male: 7.75 – 9.82 |
| **Average colour** | **Form** | Spring: 55.1 ± 7.8<br>Summer: 44.8 ± 6.9 | -3.63 | **0.001** | −10.30 Summer < Spring) | Spring: 51.1 – 59.2<br>Summer: 41.3 – 48.4 |
| | **Sex** | Female: 55.7 ± 7.6<br>Male: 44.3 ± 8.4 | -3.94 | **<0.001** | −11.38 (Male < Female) | Female: 52.3 – 59-1<br>Male: 40.1 – 48.5 |

**Appendix S4:** Individual butterfly heating and cooling curves for the thoracic and wing regions depending on status (alive, dead) and seasonal form (spring, summer)

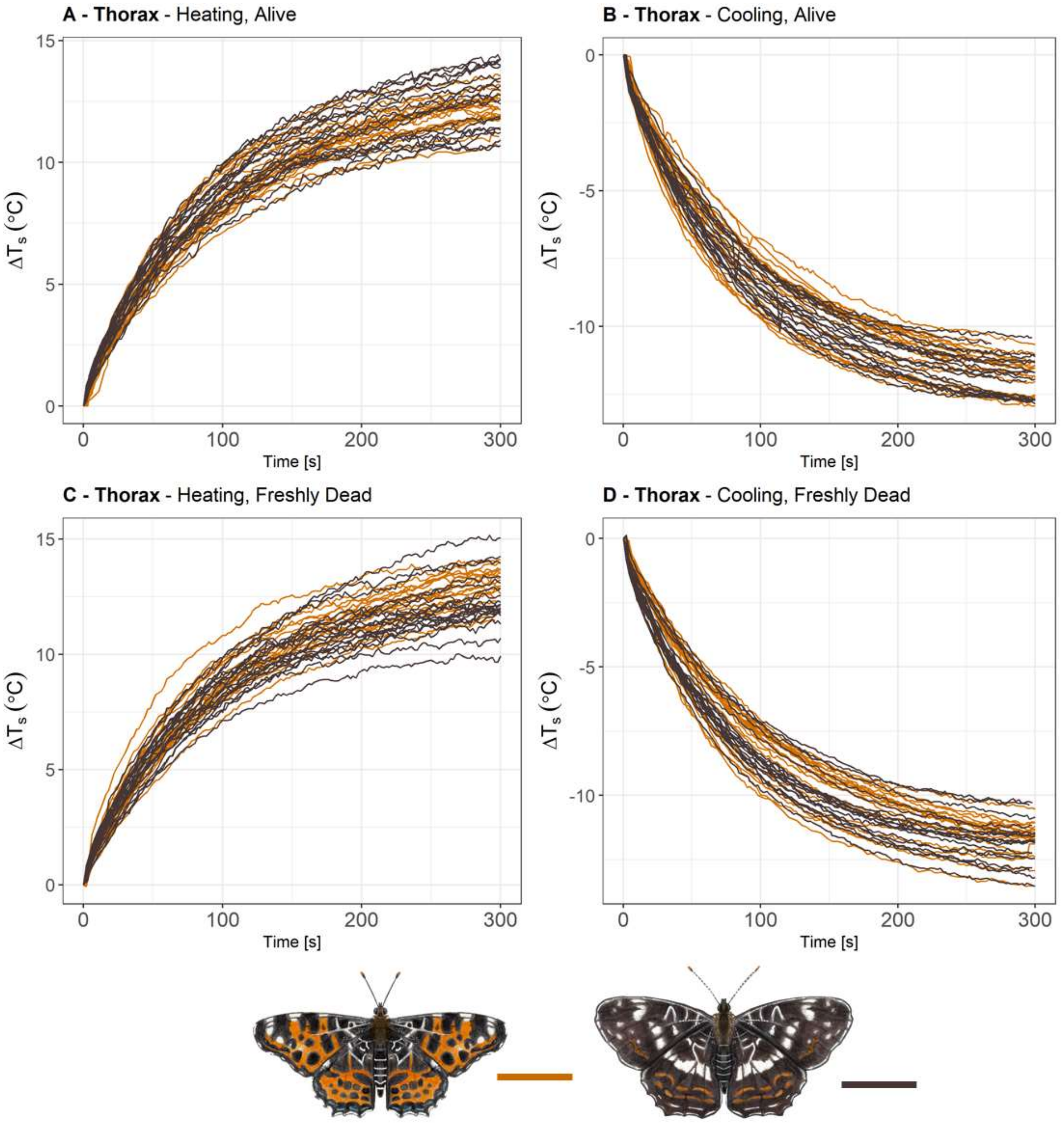


**Figure S1: Thorax** heating and cooling curves in seasonally polymorphic *Araschnia levana* butterfly. Temperature in the figure was standardised using ΔT compared to $T_0$, separated by heating and cooling by status (alive vs. freshly dead).

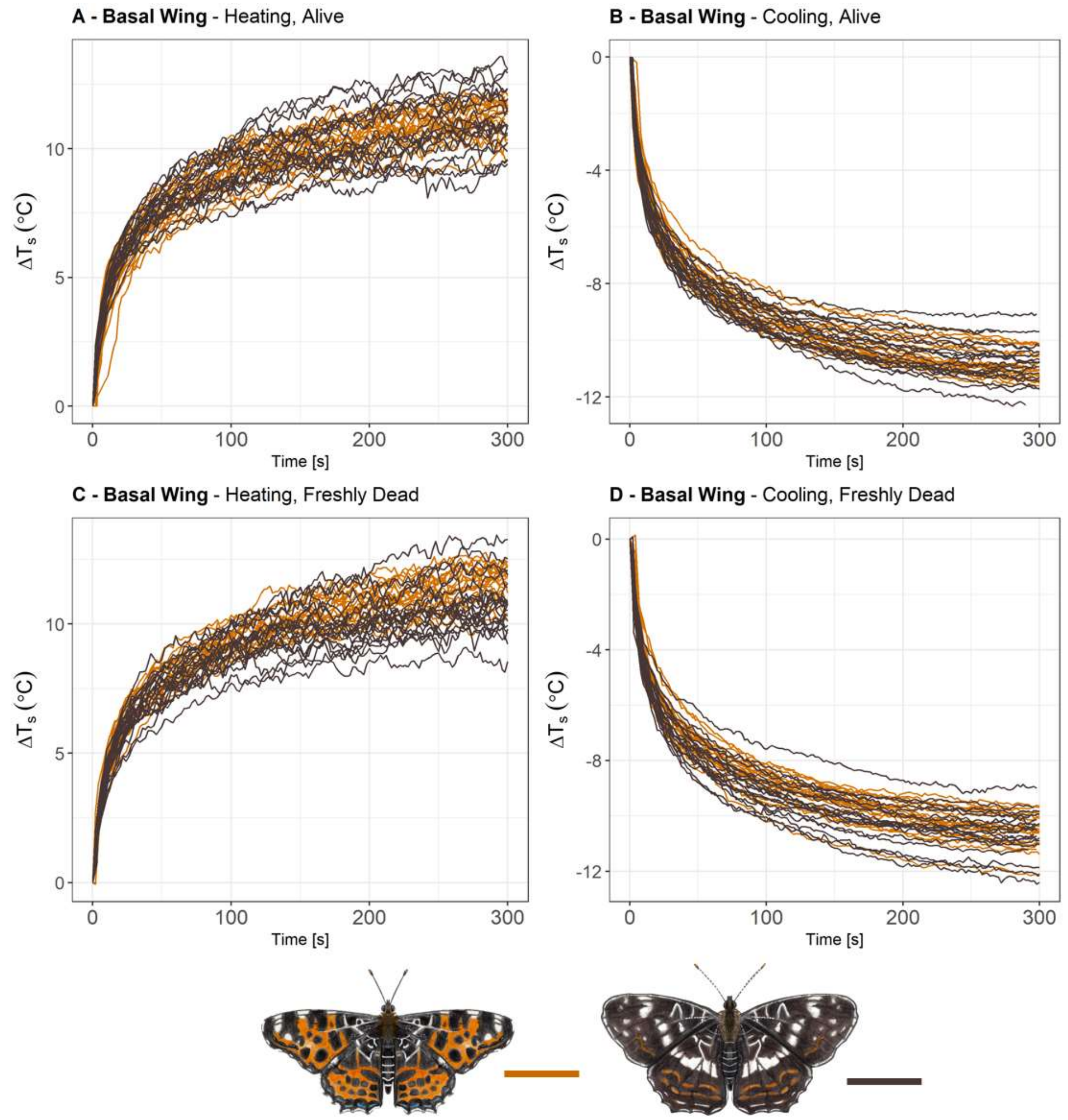


**Figure S2: Basal wing** heating and cooling curves in seasonally polymorphic *Araschnia levana* butterfly. Temperature in the figure was standardised using ΔT compared to $T_0$, separated by heating and cooling by status (alive vs. freshly dead).

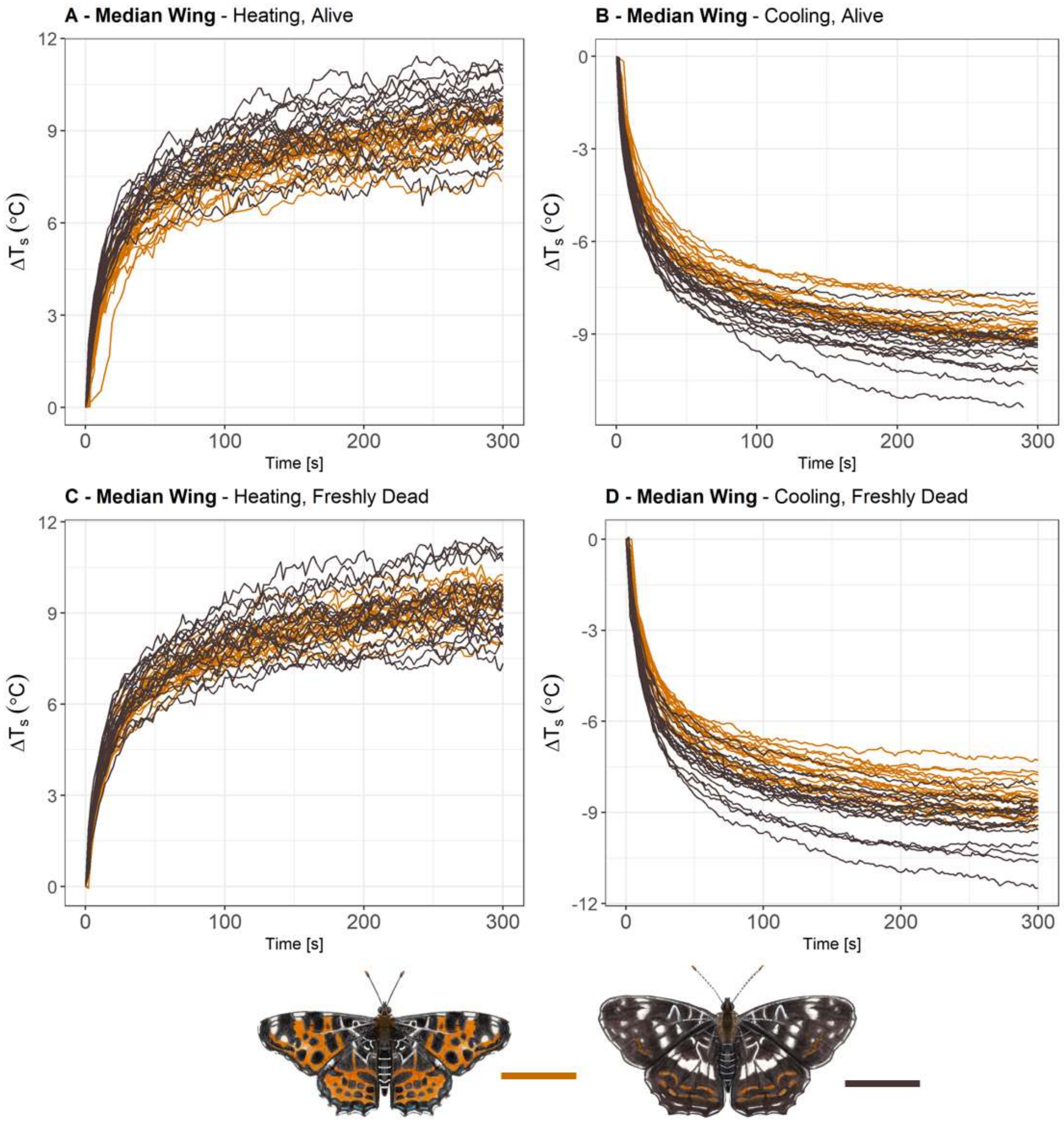


**Figure S3: Median wing** heating and cooling curves in seasonally polymorphic *Araschnia levana* butterfly. Temperature in the figure was standardised using ΔT compared to $T_0$, separated by heating and cooling by status (alive vs. freshly dead).

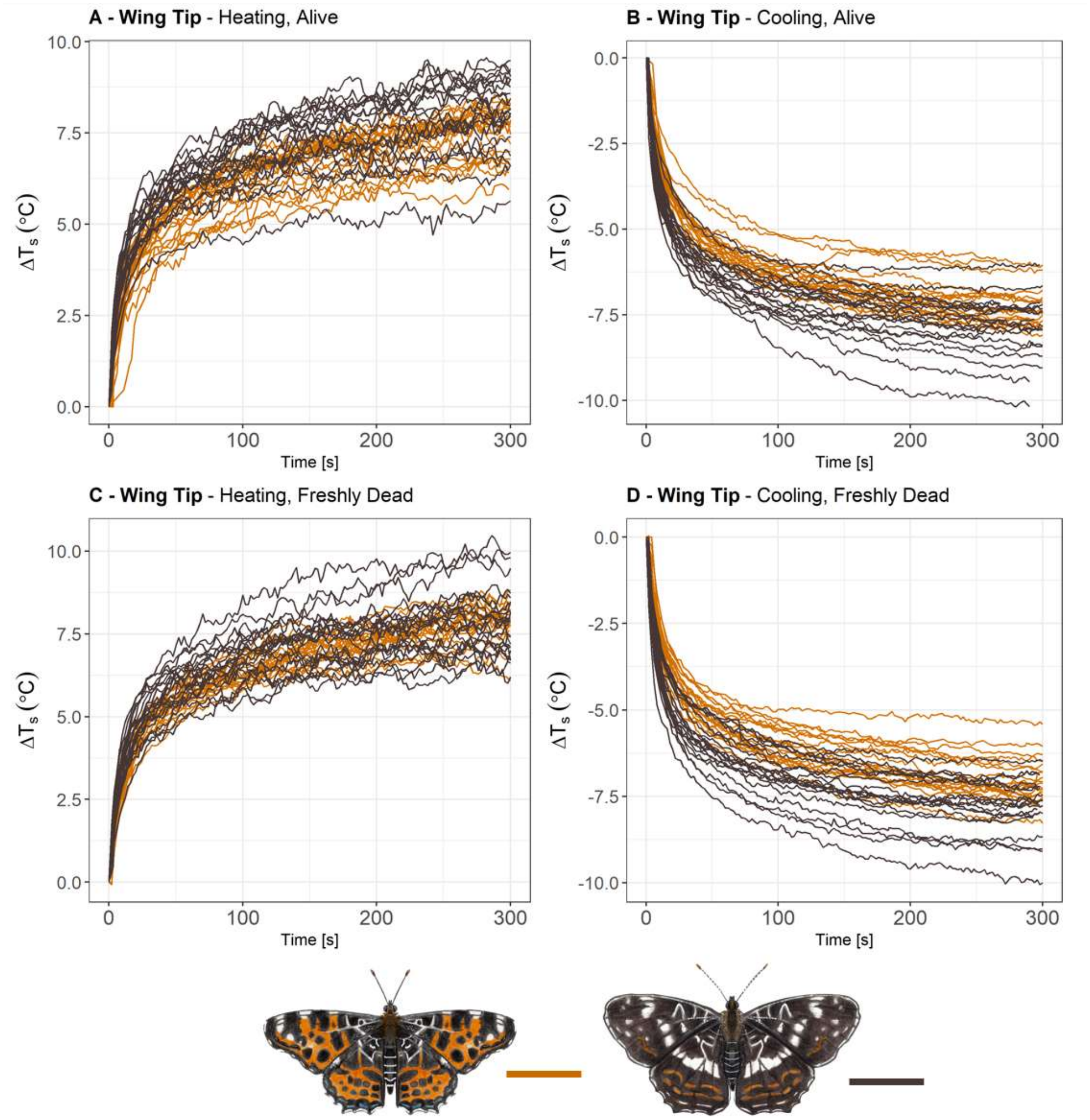


**Figure S4: Wing tip** heating and cooling curves in seasonally polymorphic *Araschnia levana* butterfly. Temperature in the figure was standardised using ΔT compared to $T_0$, separated by heating and cooling by status (alive vs. freshly dead).

**Appendix S5:** Results of the two linear mixed effect models (one using wing loading the other using wing area and weight) for each ROI (heating and cooling) including contrasts at each timeframe for categorical parameters.

**Table S3-1: Thorax heating**: Output of the **wing loading** model and contrasts tables for status (alive vs. freshly dead) and form (spring vs. summer), all continuous factors were scaled to assure equal variance. Additional contrast tables for continuous factors are not included but can be created using the provided scripts.

| Term | Sum Sq | Mean Sq | NumDF | DenDF | F value | Pr(>F) | Signif |
|---|---|---|---|---|---|---|---|
| Average colour | 0.1 | 0.06 | 1 | 35.654 | 0.3432 | 0.56168 | |
| Wing loading | 4.2 | 4.17 | 1 | 35.437 | 22.2225 | **3.70E-05** | *** |
| Form | 0.6 | 0.62 | 1 | 39.665 | 3.29 | 0.07727 | . |
| Sex | 0 | 0.03 | 1 | 36.734 | 0.1587 | 0.69268 | |
| Status | 0.1 | 0.06 | 1 | 39.493 | 0.3462 | 0.55961 | |
| Time (factor) | 6727.1 | 1681.79 | 4 | 298.256 | 8973.101 | **<2.2e-16** | *** |
| Average colour:Time | 0 | 0.01 | 4 | 298.256 | 0.0591 | 0.99351 | |
| Wing loading:Time | 5 | 1.24 | 4 | 298.22 | 6.6189 | **4.08E-05** | *** |
| Form:Time | 1.9 | 0.49 | 4 | 298.225 | 2.5888 | **0.03698** | * |
| Status:Time | 0.2 | 0.05 | 4 | 298.245 | 0.2813 | 0.88998 | |

**Table S3-2:** Contrasts for **status** (alive vs. freshly dead) depending on time.

| Time | Contrast | Estimate | SE | Df | t-ratio | p value |
|---|---|---|---|---|---|---|
| t10 | Alive – Freshly dead | -0.0358 | 0.14 | 97.5 | -0.255 | 0.7989 |
| t30 | Alive – Freshly dead | -0.0296 | 0.14 | 97.5 | -0.211 | 0.8336 |
| t60 | Alive – Freshly dead | -0.034 | 0.14 | 97.5 | -0.242 | 0.809 |
| t180 | Alive – Freshly dead | -0.0727 | 0.14 | 97.5 | -0.518 | 0.6057 |
| t300 | Alive – Freshly dead | -0.1526 | 0.141 | 100.2 | -1.079 | 0.2831 |

**Table S3-3:** Contrasts for **form** (spring vs. summer) depending on time.

| Time | Contrast | Estimate | SE | Df | t-ratio | p value |
|---|---|---|---|---|---|---|
| t10 | spring - summer | -0.0468 | 0.236 | 82.8 | -0.198 | 0.8432 |
| t30 | spring - summer | 0.3186 | 0.236 | 82.8 | 1.35 | 0.1807 |
| t60 | spring - summer | 0.5008 | 0.236 | 82.8 | 2.122 | **0.0368** |
| t180 | spring - summer | 0.5519 | 0.236 | 82.8 | 2.339 | **0.0218** |
| t300 | spring - summer | 0.4389 | 0.237 | 84.5 | 1.849 | 0.0679 |

**Table S4-1: Thorax heating**: Output of the **area and weight** model and contrasts tables for status (alive vs. freshly dead) and form (spring vs. summer), all continuous factors were scaled to assure equal variance. Additional contrast tables for continuous factors are not included but can be created using the provided scripts.

| Term | Sum Sq | Mean Sq | NumDF | DenDF | F value | Pr(>F) | Signif |
|---|---|---|---|---|---|---|---|
| Average colour | 0 | 0.04 | 1 | 34.122 | 0.2007 | 0.657015 | |
| Weight | 3.6 | 3.63 | 1 | 33.946 | 20.1005 | **7.97E-05** | *** |
| Total wing area | 1.4 | 1.36 | 1 | 35.873 | 7.5314 | **0.009409** | ** |
| Form | 0.7 | 0.72 | 1 | 39.471 | 3.9621 | 0.053493 | . |
| Sex | 0 | 0.04 | 1 | 36.306 | 0.203 | 0.654995 | |
| Status | 0.1 | 0.06 | 1 | 39.209 | 0.3148 | 0.577926 | |
| Time (factor) | 6727.4 | 1681.85 | 4 | 294.253 | 9309.781 | **<2.2e-16** | *** |
| Average colour:Time | 0.2 | 0.05 | 4 | 294.281 | 0.2575 | 0.904959 | |
| Weight:Time | 5.1 | 1.27 | 4 | 294.212 | 7.0533 | **1.96E-05** | *** |
| Total wing area:Time | 2.6 | 0.65 | 4 | 294.24 | 3.5717 | **0.007302** | ** |
| Form:Time | 2.6 | 0.66 | 4 | 294.258 | 3.6405 | **0.006506** | ** |
| Status:Time | 0.2 | 0.04 | 4 | 294.241 | 0.2201 | 0.927103 | |

**Table S4-2:** Contrasts for **status** (alive vs. freshly dead) depending on time.

| Time | Contrast | Estimate | SE | Df | t-ratio | p value |
|---|---|---|---|---|---|---|
| t10 | Alive – Freshly dead | -0.0353 | 0.14 | 94.8 | -0.253 | 0.8007 |
| t30 | Alive – Freshly dead | -0.0315 | 0.14 | 94.8 | -0.226 | 0.8218 |
| t60 | Alive – Freshly dead | -0.036 | 0.14 | 94.8 | -0.258 | 0.7968 |
| t180 | Alive – Freshly dead | -0.0694 | 0.14 | 94.8 | -0.497 | 0.6201 |
| t300 | Alive – Freshly dead | -0.1381 | 0.141 | 97.4 | -0.981 | 0.3289 |

**Table S4-3:** Contrasts for **form** (spring vs. summer) depending on time.

| Time | Contrast | Estimate | SE | Df | t-ratio | p value |
|---|---|---|---|---|---|---|
| t10 | spring - summer | 0.031 | 0.282 | 69.3 | 0.11 | 0.9128 |
| t30 | spring - summer | 0.351 | 0.282 | 69.3 | 1.246 | 0.2168 |
| t60 | spring - summer | 0.55 | 0.282 | 69.3 | 1.953 | 0.0549 |
| t180 | spring - summer | 0.743 | 0.282 | 69.3 | 2.639 | **0.0103** |
| t300 | spring - summer | 0.747 | 0.283 | 71.1 | 2.634 | **0.0104** |

**Table S5-1: Thorax cooling**: Output of the **wing loading** model and contrasts tables for status (alive vs. freshly dead) and form (spring vs. summer), all continuous factors were scaled to assure equal variance. Additional contrast tables for continuous factors are not included but can be created using the provided scripts.

| Term | Sum Sq | Mean Sq | NumDF | DenDF | F value | Pr(>F) | Signif |
|---|---|---|---|---|---|---|---|
| Average colour | 0.2 | 0.15 | 1 | 33.395 | 1.4259 | 0.2408559 | |
| Wing loading | 4.6 | 4.61 | 1 | 33.661 | 43.4643 | **1.551E-07** | *** |
| Form | 0.5 | 0.54 | 1 | 36.285 | 5.0936 | **0.0301303** | * |
| Sex | 0 | 0.04 | 1 | 34.007 | 0.351 | 0.5574528 | |
| Status | 0.1 | 0.12 | 1 | 35.529 | 1.1493 | 0.2909216 | |
| Time (factor) | 5621.4 | 1405.34 | 4 | 280.793 | 13249.73 | **<2.2e-16** | *** |
| Average colour:Time | 2 | 0.51 | 4 | 280.635 | 4.7724 | **0.0009703** | ** |
| Wing loading:Time | 8 | 2.01 | 4 | 280.457 | 18.9417 | **8.418E-14** | *** |
| Form:Time | 2.8 | 0.71 | 4 | 280.459 | 6.6795 | **3.791E-05** | *** |
| Status:Time | 0.1 | 0.02 | 4 | 280.787 | 0.2292 | 0.921867 | |

**Table S5-2:** Contrasts for **status** (alive vs. freshly dead) depending on time.

| Time | Contrast | Estimate | SE | Df | t-ratio | p value |
|---|---|---|---|---|---|---|
| t10 | Alive – Freshly dead | -0.0418 | 0.0917 | 140 | -0.456 | 0.649 |
| t30 | Alive – Freshly dead | -0.0489 | 0.0917 | 140 | -0.534 | 0.5945 |
| t60 | Alive – Freshly dead | -0.0551 | 0.0922 | 143 | -0.598 | 0.5511 |
| t180 | Alive – Freshly dead | -0.0609 | 0.0922 | 142 | -0.661 | 0.5099 |
| t300 | Alive – Freshly dead | -0.1329 | 0.0949 | 154 | -1.4 | 0.1634 |

**Table S5-3:** Contrasts for **form** (spring vs. summer) depending on time.

| Time | Contrast | Estimate | SE | Df | t-ratio | p value |
|---|---|---|---|---|---|---|
| t10 | spring - summer | 0.0519 | 0.195 | 68.7 | 0.265 | 0.7915 |
| t30 | spring - summer | -0.1982 | 0.195 | 68.7 | -1.014 | 0.3141 |
| t60 | spring - summer | -0.5033 | 0.196 | 69 | -2.573 | **0.0122** |
| t180 | spring - summer | -0.6256 | 0.195 | 68.7 | -3.202 | **0.0021** |
| t300 | spring - summer | -0.6083 | 0.197 | 71.3 | -3.082 | **0.0029** |

**Table S6-1: Thorax cooling**: Output of the **area and weight** model and contrasts tables for status (alive vs. freshly dead) and form (spring vs. summer), all continuous factors were scaled to assure equal variance. Additional contrast tables for continuous factors are not included but can be created using the provided scripts.

| Term | Sum Sq | Mean Sq | NumDF | DenDF | F value | Pr(>F) | Signif |
|---|---|---|---|---|---|---|---|
| Average colour | 0.1 | 0.15 | 1 | 32.247 | 1.4188 | 0.2422987 | |
| Weight | 4.5 | 4.49 | 1 | 32.403 | 42.9859 | **2.064E-07** | *** |
| Total wing area | 0.8 | 0.76 | 1 | 33.693 | 7.2616 | **0.0109051** | * |
| Form | 0.3 | 0.34 | 1 | 36.15 | 3.2389 | 0.0802608 | . |
| Sex | 0 | 0.01 | 1 | 33.568 | 0.1316 | 0.7190121 | |
| Status | 0.1 | 0.12 | 1 | 35.514 | 1.1018 | 0.3009675 | |
| Time (factor) | 5618.6 | 1404.65 | 4 | 276.791 | 13450.94 | **<2.2e-16** | *** |
| Average colour:Time | 2.2 | 0.56 | 4 | 276.589 | 5.3326 | **0.0003763** | ** |
| Weight:Time | 8.3 | 2.07 | 4 | 276.496 | 19.814 | **2.344E-14** | *** |
| Total wing area:Time | 1.5 | 0.38 | 4 | 276.756 | 3.6549 | **0.0064035** | ** |
| Form:Time | 2.5 | 0.62 | 4 | 276.74 | 5.9646 | **0.0001285** | *** |
| Status:Time | 0.1 | 0.03 | 4 | 276.802 | 0.2459 | 0.911988 | |

**Table S6-2:** Contrasts for **status** (alive vs. freshly dead) depending on time.

| Time | Contrast | Estimate | SE | Df | t-ratio | p value |
|---|---|---|---|---|---|---|
| t10 | Alive – Freshly dead | -0.0439 | 0.0911 | 139 | -0.482 | 0.6305 |
| t30 | Alive – Freshly dead | -0.0462 | 0.0911 | 139 | -0.507 | 0.613 |
| t60 | Alive – Freshly dead | -0.0485 | 0.0916 | 142 | -0.53 | 0.5971 |
| t180 | Alive – Freshly dead | -0.0587 | 0.0917 | 142 | -0.64 | 0.5229 |
| t300 | Alive – Freshly dead | -0.1335 | 0.0943 | 153 | -1.416 | 0.1588 |

**Table S6-3:** Contrasts for **form** (spring vs. summer) depending on time.

| Time | Contrast | Estimate | SE | Df | t-ratio | p value |
|---|---|---|---|---|---|---|
| t10 | spring - summer | 0.0163 | 0.237 | 59.2 | 0.069 | 0.9452 |
| t30 | spring - summer | -0.1579 | 0.237 | 59.2 | -0.667 | 0.5076 |
| t60 | spring - summer | -0.4391 | 0.237 | 59.6 | -1.85 | 0.0692 |
| t180 | spring - summer | -0.6125 | 0.237 | 59.4 | -2.583 | **0.0123** |
| t300 | spring - summer | -0.6983 | 0.24 | 62.2 | -2.91 | **0.005** |

**Table S7-1: Wing tip heating**: Output of the **wing loading** model and contrasts tables for status (alive vs. freshly dead) and form (spring vs. summer), all continuous factors were scaled to assure equal variance. Additional contrast tables for continuous factors are not included but can be created using the provided scripts.

| Term | Sum Sq | Mean Sq | NumDF | DenDF | F value | Pr(>F) | Signif |
|---|---|---|---|---|---|---|---|
| Average colour | 1.41 | 1.413 | 1 | 36.253 | 9.8947 | **0.003304** | ** |
| Wing loading | 0.05 | 0.052 | 1 | 36.042 | 0.3616 | 0.551374 | |
| Form | 0.48 | 0.479 | 1 | 40.139 | 3.3541 | 0.074467 | . |
| Sex | 0.91 | 0.907 | 1 | 37.306 | 6.3486 | **0.016156** | * |
| Status | 0.03 | 0.03 | 1 | 40.073 | 0.2066 | 0.651867 | |
| Time (factor) | 1188.27 | 297.067 | 4 | 298.174 | 2080.292 | **<2.2e-16** | *** |
| Average colour:Time | 0.73 | 0.182 | 4 | 298.173 | 1.2747 | 0.279926 | |
| Wing loading:Time | 0.2 | 0.05 | 4 | 298.145 | 0.3482 | 0.845204 | |
| Form:Time | 0.31 | 0.077 | 4 | 298.149 | 0.5396 | 0.706767 | |
| Status:Time | 0.06 | 0.014 | 4 | 298.165 | 0.0986 | 0.982845 | |

**Table S7-2:** Contrasts for **status** (alive vs. freshly dead) depending on time.

| Time | Contrast | Estimate | SE | Df | t-ratio | p value |
|---|---|---|---|---|---|---|
| t10 | Alive – Freshly dead | -0.0509 | 0.13 | 86.6 | -0.392 | 0.696 |
| t30 | Alive – Freshly dead | -0.0622 | 0.13 | 86.6 | -0.479 | 0.6333 |
| t60 | Alive – Freshly dead | -0.0797 | 0.13 | 86.6 | -0.614 | 0.5408 |
| t180 | Alive – Freshly dead | -0.0387 | 0.13 | 86.6 | -0.298 | 0.7666 |
| t300 | Alive – Freshly dead | -0.0084 | 0.131 | 88.8 | -0.065 | 0.9487 |

**Table S7-3:** Contrasts for **form** (spring vs. summer) depending on time.

| Time | Contrast | Estimate | SE | Df | t-ratio | p value |
|---|---|---|---|---|---|---|
| t10 | spring - summer | -0.503 | 0.227 | 71.3 | -2.216 | **0.0299** |
| t30 | spring - summer | -0.328 | 0.227 | 71.3 | -1.446 | 0.1525 |
| t60 | spring - summer | -0.371 | 0.227 | 71.3 | -1.636 | 0.1063 |
| t180 | spring - summer | -0.342 | 0.227 | 71.3 | -1.506 | 0.1364 |
| t300 | spring - summer | -0.237 | 0.228 | 72.6 | -1.041 | 0.3012 |

**Table S8-1: Wing tip heating**: Output of the **area and weight** model and contrasts tables for status (alive vs. freshly dead) and form (spring vs. summer), all continuous factors were scaled to assure equal variance. Additional contrast tables for continuous factors are not included but can be created using the provided scripts.

| Term | Sum Sq | Mean Sq | NumDF | DenDF | F value | Pr(>F) | Signif |
|---|---|---|---|---|---|---|---|
| Average colour | 1.22 | 1.218 | 1 | 34.992 | 8.6772 | **0.005697** | ** |
| Weight | 0.12 | 0.119 | 1 | 34.817 | 0.8475 | 0.363588 | |
| Total wing area | 0.36 | 0.357 | 1 | 36.727 | 2.5415 | 0.119461 | |
| Form | 0.01 | 0.012 | 1 | 40.281 | 0.0849 | 0.772233 | |
| Sex | 0 | 0.002 | 1 | 37.158 | 0.0155 | 0.901505 | |
| Status | 0.02 | 0.022 | 1 | 40.104 | 0.1567 | 0.694338 | |
| Time (factor) | 1189.54 | 297.385 | 4 | 294.19 | 2118.788 | **<2.2e-16** | *** |
| Average colour:Time | 0.35 | 0.088 | 4 | 294.214 | 0.6259 | 0.644393 | |
| Weight:Time | 0.25 | 0.063 | 4 | 294.155 | 0.4483 | 0.773578 | |
| Total wing area:Time | 1.07 | 0.269 | 4 | 294.179 | 1.9135 | 0.108155 | |
| Form:Time | 0.83 | 0.207 | 4 | 294.194 | 1.4783 | 0.208715 | |
| Status:Time | 0.07 | 0.017 | 4 | 294.18 | 0.1213 | 0.974824 | |

**Table S8-2:** Contrasts for **status** (alive vs. freshly dead) depending on time.

| Time | Contrast | Estimate | SE | Df | t-ratio | p value |
|---|---|---|---|---|---|---|
| t10 | Alive – Freshly dead | -0.0491 | 0.129 | 85.8 | -0.379 | 0.7056 |
| t30 | Alive – Freshly dead | -0.058 | 0.129 | 85.8 | -0.448 | 0.6551 |
| t60 | Alive – Freshly dead | -0.0742 | 0.129 | 85.8 | -0.574 | 0.5678 |
| t180 | Alive – Freshly dead | -0.03 | 0.129 | 85.8 | -0.232 | 0.8173 |
| t300 | Alive – Freshly dead | 0.0025 | 0.13 | 88.1 | 0.019 | 0.9845 |

**Table S8-3:** Contrasts for **form** (spring vs. summer) depending on time.

| Time | Contrast | Estimate | SE | Df | t-ratio | p value |
|---|---|---|---|---|---|---|
| t10 | spring - summer | -0.326 | 0.266 | 63.9 | -1.224 | 0.2253 |
| t30 | spring - summer | -0.085 | 0.266 | 63.9 | -0.318 | 0.7513 |
| t60 | spring - summer | -0.091 | 0.266 | 63.9 | -0.341 | 0.7344 |
| t180 | spring - summer | 0.032 | 0.266 | 63.9 | 0.122 | 0.9036 |
| t300 | spring - summer | 0.127 | 0.268 | 65.3 | 0.473 | 0.6376 |

**Table S9-1: Wing tip cooling**: Output of the **wing loading** model and contrasts tables for status (alive vs. freshly dead) and form (spring vs. summer), all continuous factors were scaled to assure equal variance. Additional contrast tables for continuous factors are not included but can be created using the provided scripts.

| Term | Sum Sq | Mean Sq | NumDF | DenDF | F value | Pr(>F) | Signif |
|---|---|---|---|---|---|---|---|
| Average colour | 1.49 | 1.491 | 1 | 34.359 | 12.1946 | **0.001338** | ** |
| Wing loading | 0.08 | 0.083 | 1 | 34.657 | 0.6769 | 0.416284 | |
| Form | 1.11 | 1.113 | 1 | 37.512 | 9.1085 | **0.004555** | ** |
| Sex | 0.42 | 0.423 | 1 | 35.031 | 3.4633 | 0.071156 | . |
| Status | 0.31 | 0.305 | 1 | 37.337 | 2.4969 | 0.122506 | |
| Time (factor) | 902.31 | 225.577 | 4 | 280.675 | 1845.353 | **<2.2e-16** | *** |
| Average colour:Time | 0.81 | 0.203 | 4 | 280.563 | 1.6577 | 0.160002 | |
| Wing loading:Time | 1.01 | 0.251 | 4 | 280.43 | 2.0562 | 0.086748 | . |
| Form:Time | 0.89 | 0.223 | 4 | 280.431 | 1.8281 | 0.12352 | |
| Status:Time | 0.14 | 0.034 | 4 | 280.673 | 0.2802 | 0.890672 | |

**Table S9-2:** Contrasts for **status** (alive vs. freshly dead) depending on time.

| Time | Contrast | Estimate | SE | Df | t-ratio | p value |
|---|---|---|---|---|---|---|
| t10 | Alive – Freshly dead | -0.117 | 0.109 | 106 | -1.071 | 0.2865 |
| t30 | Alive – Freshly dead | -0.071 | 0.109 | 106 | -0.648 | 0.5183 |
| t60 | Alive – Freshly dead | -0.132 | 0.109 | 107 | -1.207 | 0.2301 |
| t180 | Alive – Freshly dead | -0.141 | 0.109 | 107 | -1.286 | 0.2011 |
| t300 | Alive – Freshly dead | -0.191 | 0.112 | 116 | -1.702 | 0.0914 |

**Table S9-3:** Contrasts for **form** (spring vs. summer) depending on time.

| Time | Contrast | Estimate | SE | Df | t-ratio | p value |
|---|---|---|---|---|---|---|
| t10 | spring - summer | 0.643 | 0.231 | 61.5 | 2.789 | **0.007** |
| t30 | spring - summer | 0.806 | 0.231 | 61.5 | 3.494 | **0.0009** |
| t60 | spring - summer | 0.707 | 0.231 | 61.8 | 3.064 | **0.0032** |
| t180 | spring - summer | 0.541 | 0.231 | 61.5 | 2.344 | **0.0223** |
| t300 | spring - summer | 0.368 | 0.233 | 63.5 | 1.582 | 0.1185 |

**Table S10-1: Wing tip cooling**: Output of the **area and weight** model and contrasts tables for status (alive vs. freshly dead) and form (spring vs. summer), all continuous factors were scaled to assure equal variance. Additional contrast tables for continuous factors are not included but can be created using the provided scripts.

| Term | Sum Sq | Mean Sq | NumDF | DenDF | F value | Pr(>F) | Signif |
|---|---|---|---|---|---|---|---|
| Average colour | 1.38 | 1.375 | 1 | 32.875 | 11.2916 | **0.001985** | ** |
| Weight | 0.13 | 0.131 | 1 | 33.058 | 1.079 | 0.306459 | |
| Total wing area | 0.14 | 0.141 | 1 | 34.49 | 1.1593 | 0.289085 | |
| Form | 0.27 | 0.268 | 1 | 37.258 | 2.1977 | 0.146623 | |
| Sex | 0 | 0 | 1 | 34.373 | 0.0008 | 0.97707 | |
| Status | 0.33 | 0.326 | 1 | 37.023 | 2.6781 | 0.110211 | |
| Time (factor) | 902.68 | 225.671 | 4 | 276.691 | 1853.127 | **<2.2e-16** | *** |
| Average colour:Time | 1.1 | 0.276 | 4 | 276.543 | 2.2649 | 0.062454 | . |
| Weight:Time | 1 | 0.249 | 4 | 276.474 | 2.0436 | 0.088518 | . |
| Total wing area:Time | 0.23 | 0.058 | 4 | 276.668 | 0.4727 | 0.755762 | |
| Form:Time | 1.16 | 0.291 | 4 | 276.652 | 2.3887 | 0.051261 | . |
| Status:Time | 0.15 | 0.038 | 4 | 276.703 | 0.3082 | 0.872391 | |

**Table S10-2:** Contrasts for **status** (alive vs. freshly dead) depending on time.

| Time | Contrast | Estimate | SE | Df | t-ratio | p value |
|---|---|---|---|---|---|---|
| t10 | Alive – Freshly dead | -0.116 | 0.109 | 105 | -1.069 | 0.2877 |
| t30 | Alive – Freshly dead | -0.073 | 0.109 | 105 | -0.672 | 0.5029 |
| t60 | Alive – Freshly dead | -0.138 | 0.109 | 107 | -1.264 | 0.2091 |
| t180 | Alive – Freshly dead | -0.15 | 0.109 | 107 | -1.37 | 0.1737 |
| t300 | Alive – Freshly dead | -0.197 | 0.112 | 115 | -1.757 | 0.0816 |

**Table S10-3:** Contrasts for **form** (spring vs. summer) depending on time.

| Time | Contrast | Estimate | SE | Df | t-ratio | p value |
|---|---|---|---|---|---|---|
| t10 | spring - summer | 0.487 | 0.278 | 55.4 | 1.752 | 0.0853 |
| t30 | spring - summer | 0.604 | 0.278 | 55.4 | 2.173 | **0.0341** |
| t60 | spring - summer | 0.446 | 0.279 | 55.7 | 1.6 | 0.1153 |
| t180 | spring - summer | 0.253 | 0.278 | 55.6 | 0.91 | 0.3669 |
| t300 | spring - summer | 0.075 | 0.281 | 57.8 | 0.268 | 0.7896 |

**Table S11-1: Basal wing heating**: Output of the **wing loading** model and contrasts tables for status (alive vs. freshly dead) and form (spring vs. summer), all continuous factors were scaled to assure equal variance. Additional contrast tables for continuous factors are not included but can be created using the provided scripts.

| Term | Sum Sq | Mean Sq | NumDF | DenDF | F value | Pr(>F) | Signif |
|---|---|---|---|---|---|---|---|
| Average colour | 0.24 | 0.24 | 1 | 36.07 | 1.4793 | 0.23179 | |
| Wing loading | 0 | 0 | 1 | 35.87 | 0.0002 | 0.9888 | |
| Form | 0 | 0 | 1 | 39.69 | 0.0055 | 0.94143 | |
| Sex | 1.1 | 1.1 | 1 | 37.05 | 6.7084 | **0.01364** | * |
| Status | 0.01 | 0.01 | 1 | 39.68 | 0.0809 | 0.77754 | |
| Time (factor) | 2701.92 | 675.48 | 4 | 298.14 | 4131.456 | < **2e-16** | *** |
| Average colour:Time | 0.93 | 0.23 | 4 | 298.14 | 1.4162 | 0.22851 | |
| Wing loading:Time | 0.28 | 0.07 | 4 | 298.11 | 0.4346 | 0.78363 | |
| Form:Time | 0.83 | 0.21 | 4 | 298.11 | 1.2675 | 0.2828 | |
| Status:Time | 0.47 | 0.12 | 4 | 298.13 | 0.7183 | 0.57997 | |

**Table S11-2:** Contrasts for **status** (alive vs. freshly dead) depending on time.

| Time | Contrast | Estimate | SE | Df | t-ratio | p value |
|---|---|---|---|---|---|---|
| t10 | Alive – Freshly dead | −0.045 | 0.143 | 82.3 | −0.315 | 0.7539 |
| t30 | Alive – Freshly dead | −0.044 | 0.143 | 82.3 | −0.306 | 0.76 |
| t60 | Alive – Freshly dead | 0.037 | 0.143 | 82.3 | 0.261 | 0.9158 |
| t180 | Alive – Freshly dead | 0.104 | 0.143 | 82.3 | 0.727 | 0.4691 |
| t300 | Alive – Freshly dead | 0.115 | 0.144 | 84.3 | 0.799 | 0.4267 |

**Table S11-3:** Contrasts for **form** (spring vs. summer) depending on time.

| Time | Contrast | Estimate | SE | Df | t-ratio | p value |
|---|---|---|---|---|---|---|
| t10 | spring - summer | −0.203 | 0.268 | 63.1 | −0.756 | 0.4525 |
| t30 | spring - summer | 0.002 | 0.268 | 63.1 | 0.006 | 0.9949 |
| t60 | spring - summer | −0.028 | 0.268 | 63.1 | −0.106 | 0.9158 |
| t180 | spring - summer | 0.093 | 0.268 | 63.1 | 0.345 | 0.7309 |
| t300 | spring - summer | 0.224 | 0.269 | 64 | 0.834 | 0.4074 |

**Table S12-1: Basal wing heating**: Output of the **area and weight** model and contrasts tables for status (alive vs. freshly dead) and form (spring vs. summer), all continuous factors were scaled to assure equal variance. Additional contrast tables for continuous factors are not included but can be created using the provided scripts.

| Term | Sum Sq | Mean Sq | NumDF | DenDF | F value | Pr(>F) | Signif |
|---|---|---|---|---|---|---|---|
| Average colour | 0.13 | 0.13 | 1 | 34.56 | 0.8332 | 0.3677 | |
| Weight | 0.04 | 0.04 | 1 | 34.4 | 0.275 | 0.6034 | |
| Total wing area | 0.66 | 0.66 | 1 | 36.22 | 4.1279 | **0.0496** | * |
| Form | 0.32 | 0.32 | 1 | 39.59 | 1.9837 | 0.1668 | |
| Sex | 0 | 0 | 1 | 36.62 | 0.001 | 0.9753 | |
| Status | 0.02 | 0.02 | 1 | 39.5 | 0.1189 | 0.7321 | |
| Time (factor) | 2702.99 | 675.75 | 4 | 294.15 | 4200.088 | **< 2e-16** | *** |
| Average colour:Time | 0.32 | 0.08 | 4 | 294.17 | 0.4985 | 0.7369 | |
| Weight:Time | 0.32 | 0.08 | 4 | 294.12 | 0.4933 | 0.7407 | |
| Total wing area:Time | 1.43 | 0.36 | 4 | 294.14 | 2.2196 | 0.0669 | . |
| Form:Time | 1.69 | 0.42 | 4 | 294.16 | 2.6232 | **0.035** | * |
| Status:Time | 0.51 | 0.13 | 4 | 294.14 | 0.7978 | 0.5274 | |

**Table S12-2:** Contrasts for **status** (alive vs. freshly dead) depending on time.

| Time | Contrast | Estimate | SE | Df | t-ratio | p value |
|---|---|---|---|---|---|---|
| t10 | Alive – Freshly dead | −0.043 | 0.143 | 81.5 | −0.301 | 0.7645 |
| t30 | Alive – Freshly dead | −0.038 | 0.143 | 81.5 | −0.269 | 0.7888 |
| t60 | Alive – Freshly dead | 0.044 | 0.143 | 81.5 | 0.31 | 0.7574 |
| t180 | Alive – Freshly dead | 0.113 | 0.143 | 81.5 | 0.79 | 0.4319 |
| t300 | Alive – Freshly dead | 0.128 | 0.144 | 83.5 | 0.888 | 0.3772 |

**Table S12-3:** Contrasts for **form** (spring vs. summer) depending on time.

| Time | Contrast | Estimate | SE | Df | t-ratio | p value |
|---|---|---|---|---|---|---|
| t10 | spring - summer | 0.04 | 0.311 | 58.4 | 0.13 | 0.8972 |
| t30 | spring - summer | 0.348 | 0.311 | 58.4 | 1.117 | 0.2685 |
| t60 | spring - summer | 0.366 | 0.311 | 58.4 | 1.176 | 0.2442 |
| t180 | spring - summer | 0.537 | 0.311 | 58.4 | 1.723 | 0.0902 |
| t300 | spring - summer | 0.688 | 0.313 | 59.5 | 2.2 | **0.0317** |

**Table S13-1: Basal wing cooling**: Output of the **wing loading** model and contrasts tables for status (alive vs. freshly dead) and form (spring vs. summer), all continuous factors were scaled to assure equal variance. Additional contrast tables for continuous factors are not included but can be created using the provided scripts.

| Term | Sum Sq | Mean Sq | NumDF | DenDF | F value | Pr(>F) | Signif |
|---|---|---|---|---|---|---|---|
| Average colour | 0.76 | 0.76 | 1 | 31.78 | 5.7199 | **0.02287** | * |
| Wing loading | 0.01 | 0.01 | 1 | 32.09 | 0.0835 | 0.7744 | |
| Form | 0.09 | 0.09 | 1 | 35.1 | 0.6366 | 0.4303 | |
| Sex | 1.26 | 1.26 | 1 | 32.48 | 9.4424 | **0.00427** | ** |
| Status | 0.89 | 0.89 | 1 | 34.57 | 6.6905 | **0.01407** | * |
| Time (factor) | 2084.68 | 521.17 | 4 | 280.96 | 3900.755 | **< 2.2e-16** | *** |
| Average colour:Time | 0.97 | 0.24 | 4 | 280.84 | 1.8162 | 0.1258 | |
| Wing loading:Time | 0.61 | 0.15 | 4 | 280.68 | 1.1428 | 0.3366 | |
| Form:Time | 1.27 | 0.32 | 4 | 280.69 | 2.3831 | 0.0517 | . |
| Status:Time | 0.19 | 0.05 | 4 | 280.96 | 0.3467 | 0.8462 | |

**Table S13-2:** Contrasts for **status** (alive vs. freshly dead) depending on time.

| Time | Contrast | Estimate | SE | Df | t-ratio | p value |
|---|---|---|---|---|---|---|
| t10 | Alive – Freshly dead | −0.172 | 0.11 | 116 | −1.564 | 0.1205 |
| t30 | Alive – Freshly dead | −0.149 | 0.11 | 116 | −1.357 | 0.1774 |
| t60 | Alive – Freshly dead | −0.210 | 0.11 | 118 | −1.907 | 0.0589 |
| t180 | Alive – Freshly dead | −0.237 | 0.11 | 118 | −2.154 | **0.0332** |
| t300 | Alive – Freshly dead | −0.274 | 0.113 | 128 | −2.426 | **0.0167** |

**Table S13-3:** Contrasts for **form** (spring vs. summer) depending on time.

| Time | Contrast | Estimate | SE | Df | t-ratio | p value |
|---|---|---|---|---|---|---|
| t10 | spring - summer | 0.082 | 0.223 | 67.8 | 0.367 | 0.7146 |
| t30 | spring - summer | 0.407 | 0.223 | 67.8 | 1.828 | 0.072 |
| t60 | spring - summer | 0.299 | 0.223 | 68.2 | 1.341 | 0.1845 |
| t180 | spring - summer | 0.067 | 0.223 | 67.8 | 0.301 | 0.7646 |
| t300 | spring - summer | −0.092 | 0.225 | 70.4 | −0.407 | 0.6853 |

**Table S14-1: Basal wing cooling**: Output of the **area and weight** model and contrasts tables for status (alive vs. freshly dead) and form (spring vs. summer), all continuous factors were scaled to assure equal variance. Additional contrast tables for continuous factors are not included but can be created using the provided scripts.

| Term | Sum Sq | Mean Sq | NumDF | DenDF | F value | Pr(>F) | Signif |
|---|---|---|---|---|---|---|---|
| Average colour | 0.69 | 0.69 | 1 | 28.71 | 5.1222 | **0.03136** | * |
| Weight | 0.11 | 0.11 | 1 | 28.9 | 0.8064 | 0.3766 | |
| Total wing area | 0.51 | 0.51 | 1 | 30.49 | 3.7793 | 0.06117 | . |
| Form | 0.07 | 0.07 | 1 | 33.65 | 0.4945 | 0.4867 | |
| Sex | 0.01 | 0.01 | 1 | 30.38 | 0.0583 | 0.8108 | |
| Status | 0.93 | 0.93 | 1 | 32.83 | 6.8675 | **0.01319** | * |
| Time (factor) | 2084.38 | 521.1 | 4 | 277.01 | 3859.274 | < **2e-16** | *** |
| Average colour:Time | 0.85 | 0.21 | 4 | 276.84 | 1.5653 | 0.1838 | |
| Weight:Time | 0.7 | 0.17 | 4 | 276.76 | 1.2879 | 0.2749 | |
| Total wing area:Time | 0.37 | 0.09 | 4 | 276.98 | 0.6932 | 0.5972 | |
| Form:Time | 1.16 | 0.29 | 4 | 276.96 | 2.1481 | 0.0751 | . |
| Status:Time | 0.19 | 0.05 | 4 | 277.03 | 0.3609 | 0.8363 | |

**Table S14-2:** Contrasts for **status** (alive vs. freshly dead) depending on time.

| Time | Contrast | Estimate | SE | Df | t-ratio | p value |
|---|---|---|---|---|---|---|
| t10 | Alive – Freshly dead | −0.175 | 0.111 | 115 | −1.583 | 0.1162 |
| t30 | Alive – Freshly dead | −0.154 | 0.111 | 115 | −1.386 | 0.1685 |
| t60 | Alive – Freshly dead | −0.216 | 0.111 | 117 | −1.941 | 0.0546 |
| t180 | Alive – Freshly dead | −0.244 | 0.111 | 117 | −2.191 | **0.0304** |
| t300 | Alive – Freshly dead | −0.282 | 0.114 | 126 | −2.467 | **0.015** |

**Table S14-3:** Contrasts for **form** (spring vs. summer) depending on time.

| Time | Contrast | Estimate | SE | Df | t-ratio | p value |
|---|---|---|---|---|---|---|
| t10 | spring - summer | −0.193 | 0.257 | 63.7 | −0.751 | 0.4553 |
| t30 | spring - summer | 0.111 | 0.257 | 63.7 | 0.431 | 0.6683 |
| t60 | spring - summer | −0.026 | 0.258 | 64.2 | −0.100 | 0.9207 |
| t180 | spring - summer | −0.256 | 0.257 | 64 | −0.995 | 0.3235 |
| t300 | spring - summer | −0.427 | 0.261 | 67.2 | −1.636 | 0.1065 |

**Table S15-1: Median wing heating**: Output of the **wing loading** model and contrasts tables for status (alive vs. freshly dead) and form (spring vs. summer), all continuous factors were scaled to assure equal variance. Additional contrast tables for continuous factors are not included but can be created using the provided scripts.

| Term | Sum Sq | Mean Sq | NumDF | DenDF | F value | Pr(>F) | Signif |
|---|---|---|---|---|---|---|---|
| Average colour | 0.79 | 0.79 | 1 | 35.87 | 5.3482 | **0.02659** | * |
| Wing loading | 0.22 | 0.22 | 1 | 35.65 | 1.5147 | 0.22648 | |
| Form | 0.66 | 0.66 | 1 | 40.01 | 4.4506 | **0.04119** | * |
| Sex | 0.48 | 0.48 | 1 | 36.99 | 3.278 | 0.07834 | . |
| Status | 0 | 0 | 1 | 39.92 | 0.0113 | 0.91587 | |
| Time (factor) | 1876.63 | 469.16 | 4 | 298.09 | 3185.456 | **< 2e-16** | *** |
| Average colour:Time | 0.94 | 0.24 | 4 | 298.09 | 1.6035 | 0.17334 | |
| Wing loading:Time | 0.18 | 0.05 | 4 | 298.06 | 0.3116 | 0.87011 | |
| Form:Time | 0.17 | 0.04 | 4 | 298.06 | 0.2857 | 0.88716 | |
| Status:Time | 0.22 | 0.06 | 4 | 298.08 | 0.3753 | 0.82618 | |

**Table S15-2:** Contrasts for **status** (alive vs. freshly dead) depending on time.

| Time | Contrast | Estimate | SE | Df | t-ratio | p value |
|---|---|---|---|---|---|---|
| t10 | Alive – Freshly dead | -0.053 | 0.14 | 78.4 | -0.376 | 0.7082 |
| t30 | Alive – Freshly dead | -0.03 | 0.14 | 78.4 | -0.211 | 0.8334 |
| t60 | Alive – Freshly dead | 0.02 | 0.14 | 78.4 | 0.141 | 0.8881 |
| t180 | Alive – Freshly dead | 0.054 | 0.14 | 78.4 | 0.383 | 0.7027 |
| t300 | Alive – Freshly dead | 0.071 | 0.141 | 80.2 | 0.504 | 0.616 |

**Table S15-3:** Contrasts for **form** (spring vs. summer) depending on time.

| Time | Contrast | Estimate | SE | Df | t-ratio | p value |
|---|---|---|---|---|---|---|
| t10 | spring - summer | -0.47 | 0.226 | 73.5 | -2.079 | 0.0411 |
| t30 | spring - summer | -0.382 | 0.226 | 73.5 | -1.69 | 0.0953 |
| t60 | spring - summer | -0.471 | 0.226 | 73.5 | **-2.081** | **0.0409** |
| t180 | spring - summer | -0.408 | 0.226 | 73.5 | -1.802 | 0.0756 |
| t300 | spring - summer | -0.299 | 0.227 | 74.9 | -1.318 | 0.1917 |

**Table S16-1: Median wing heating**: Output of the **area and weight** model and contrasts tables for status (alive vs. freshly dead) and form (spring vs. summer), all continuous factors were scaled to assure equal variance. Additional contrast tables for continuous factors are not included but can be created using the provided scripts.

| Term | Sum Sq | Mean Sq | NumDF | DenDF | F value | Pr(>F) | Signif |
|---|---|---|---|---|---|---|---|
| Average colour | 0.62 | 0.62 | 1 | 34.72 | 4.3769 | **0.04381** | * |
| Weight | 0.35 | 0.35 | 1 | 34.53 | 2.4374 | 0.12759 | |
| Total wing area | 0.2 | 0.2 | 1 | 36.53 | 1.4029 | 0.24389 | |
| Form | 0.05 | 0.05 | 1 | 40.29 | 0.3609 | 0.55138 | |
| Sex | 0.01 | 0.01 | 1 | 37.01 | 0.0601 | 0.80767 | |
| Status | 0 | 0 | 1 | 40.04 | 0.0269 | 0.87052 | |
| Time (factor) | 1878.93 | 469.73 | 4 | 294.11 | 3318.14 | < **2.2e-16** | *** |
| Average colour:Time | 0.26 | 0.07 | 4 | 294.13 | 0.4621 | 0.76354 | |
| Weight:Time | 0.23 | 0.06 | 4 | 294.08 | 0.4074 | 0.8033 | |
| Total wing area:Time | 2.03 | 0.51 | 4 | 294.1 | 3.5898 | **0.00708** | ** |
| Form:Time | 0.75 | 0.19 | 4 | 294.11 | 1.3322 | 0.25797 | |
| Status:Time | 0.26 | 0.07 | 4 | 294.1 | 0.4645 | 0.7618 | |

**Table S16-2:** Contrasts for **status** (alive vs. freshly dead) depending on time.

| Time | Contrast | Estimate | SE | Df | t-ratio | p value |
|---|---|---|---|---|---|---|
| t10 | Alive – Freshly dead | −0.052 | 0.139 | 76.9 | −0.371 | 0.7118 |
| t30 | Alive – Freshly dead | −0.026 | 0.139 | 76.9 | −0.188 | 0.8516 |
| t60 | Alive – Freshly dead | 0.026 | 0.139 | 76.9 | 0.185 | 0.8534 |
| t180 | Alive – Freshly dead | 0.063 | 0.139 | 76.9 | 0.454 | 0.6512 |
| t300 | Alive – Freshly dead | 0.085 | 0.14 | 78.7 | 0.605 | 0.5471 |

**Table S16-3:** Contrasts for **form** (spring vs. summer) depending on time.

| Time | Contrast | Estimate | SE | Df | t-ratio | p value |
|---|---|---|---|---|---|---|
| t10 | spring - summer | −0.348 | 0.266 | 64.5 | −1.306 | 0.196 |
| t30 | spring - summer | −0.189 | 0.266 | 64.5 | −0.710 | 0.4804 |
| t60 | spring - summer | −0.202 | 0.266 | 64.5 | −0.757 | 0.4518 |
| t180 | spring - summer | −0.041 | 0.266 | 64.5 | −0.152 | 0.8797 |
| t300 | spring - summer | 0.074 | 0.268 | 65.9 | 0.277 | 0.7829 |

**Table S17-1: Median wing cooling**: Output of the **wing loading** model and contrasts tables for status (alive vs. freshly dead) and form (spring vs. summer), all continuous factors were scaled to assure equal variance. Additional contrast tables for continuous factors are not included but can be created using the provided scripts.

| Term | Sum Sq | Mean Sq | NumDF | DenDF | F value | Pr(>F) | Signif |
|---|---|---|---|---|---|---|---|
| Average colour | 1.36 | 1.36 | 1 | 34.1 | 10.225 | **0.002987** | ** |
| Wing loading | 0.22 | 0.22 | 1 | 34.41 | 1.6579 | 0.20649 | |
| Form | 1.93 | 1.93 | 1 | 37.76 | 14.5305 | **0.000495** | *** |
| Sex | 0.47 | 0.47 | 1 | 34.86 | 3.5439 | 0.06814 | . |
| Status | 0.6 | 0.6 | 1 | 37.58 | 4.5082 | **0.04038** | * |
| Time (factor) | 1557.08 | 389.27 | 4 | 280.71 | 2924.75 | **< 2.2e-16** | *** |
| Average colour:Time | 0.76 | 0.19 | 4 | 280.56 | 1.4246 | 0.22591 | |
| Wing loading:Time | 0.59 | 0.15 | 4 | 280.4 | 1.1149 | 0.34969 | |
| Form:Time | 1.07 | 0.27 | 4 | 280.4 | 2.013 | 0.0928 | . |
| Status:Time | 0.11 | 0.03 | 4 | 280.71 | 0.1978 | 0.9394 | |

**Table S17-2:** Contrasts for **status** (alive vs. freshly dead) depending on time.

| Time | Contrast | Estimate | SE | Df | t-ratio | p value |
|---|---|---|---|---|---|---|
| t10 | Alive – Freshly dead | −0.131 | 0.107 | 123 | −1.220 | 0.2246 |
| t30 | Alive – Freshly dead | −0.125 | 0.107 | 123 | −1.161 | 0.2479 |
| t60 | Alive – Freshly dead | −0.175 | 0.108 | 125 | −1.616 | 0.1086 |
| t180 | Alive – Freshly dead | −0.177 | 0.108 | 125 | −1.637 | 0.1042 |
| t300 | Alive – Freshly dead | −0.218 | 0.111 | 135 | −1.962 | 0.0518 |

**Table S17-3:** Contrasts for **form** (spring vs. summer) depending on time.

| Time | Contrast | Estimate | SE | Df | t-ratio | p value |
|---|---|---|---|---|---|---|
| t10 | spring - summer | 0.492 | 0.202 | 78.8 | 2.436 | **0.0171** |
| t30 | spring - summer | 0.845 | 0.202 | 78.8 | 4.178 | **0.0001** |
| t60 | spring - summer | 0.796 | 0.202 | 79.3 | 3.93 | **0.0002** |
| t180 | spring - summer | 0.621 | 0.202 | 78.8 | 3.073 | **0.0029** |
| t300 | spring - summer | 0.429 | 0.205 | 82.3 | 2.095 | **0.0392** |

**Table S18-1: Median wing cooling**: Output of the **area and weight** model and contrasts tables for status (alive vs. freshly dead) and form (spring vs. summer), all continuous factors were scaled to assure equal variance. Additional contrast tables for continuous factors are not included but can be created using the provided scripts.

| Term | Sum Sq | Mean Sq | NumDF | DenDF | F value | Pr(>F) | Signif |
|---|---|---|---|---|---|---|---|
| Average colour | 1.24 | 1.24 | 1 | 32.54 | 9.3985 | **0.004346** | ** |
| Weight | 0.33 | 0.33 | 1 | 32.72 | 2.4847 | 0.124581 | |
| Total wing area | 0.12 | 0.12 | 1 | 34.39 | 0.9184 | 0.344573 | |
| Form | 0.57 | 0.57 | 1 | 37.67 | 4.3089 | **0.044793** | * |
| Sex | 0 | 0 | 1 | 34.28 | 0.0015 | 0.969455 | |
| Status | 0.63 | 0.63 | 1 | 37.21 | 4.7777 | **0.035197** | * |
| Time (factor) | 1557.44 | 389.36 | 4 | 276.72 | 2944.386 | < **2.2e-16** | *** |
| Average colour:Time | 1.13 | 0.28 | 4 | 276.53 | 2.1372 | 0.07643 | . |
| Weight:Time | 0.59 | 0.15 | 4 | 276.44 | 1.114 | 0.350142 | |
| Total wing area:Time | 0.46 | 0.11 | 4 | 276.69 | 0.8662 | 0.484668 | |
| Form:Time | 1.11 | 0.28 | 4 | 276.67 | 2.0905 | 0.082249 | . |
| Status:Time | 0.12 | 0.03 | 4 | 276.74 | 0.2319 | 0.920327 | |

**Table S18-2:** Contrasts for **status** (alive vs. freshly dead) depending on time.

| Time | Contrast | Estimate | SE | Df | t-ratio | p value |
|---|---|---|---|---|---|---|
| t10 | Alive – Freshly dead | −0.131 | 0.107 | 123 | −1.218 | 0.2255 |
| t30 | Alive – Freshly dead | −0.127 | 0.107 | 123 | −1.185 | 0.2382 |
| t60 | Alive – Freshly dead | −0.181 | 0.108 | 124 | −1.676 | 0.0962 |
| t180 | Alive – Freshly dead | −0.186 | 0.108 | 125 | −1.725 | 0.087 |
| t300 | Alive – Freshly dead | −0.224 | 0.111 | 134 | −2.025 | **0.0449** |

**Table S18-3:** Contrasts for **form** (spring vs. summer) depending on time.

| Time | Contrast | Estimate | SE | Df | t-ratio | p value |
|---|---|---|---|---|---|---|
| t10 | spring - summer | 0.377 | 0.24 | 68.5 | 1.566 | 0.1219 |
| t30 | spring - summer | 0.682 | 0.24 | 68.5 | 2.837 | **0.006** |
| t60 | spring - summer | 0.565 | 0.241 | 69.1 | 2.345 | **0.0219** |
| t180 | spring - summer | 0.366 | 0.241 | 68.8 | 1.521 | 0.1329 |
| t300 | spring - summer | 0.154 | 0.244 | 72.7 | 0.628 | 0.5318 |
